# cAMP-related second messenger pathways modulate hearing function in *Aedes aegypti* mosquitoes

**DOI:** 10.1101/2025.03.23.644695

**Authors:** YiFeng Y.J. Xu, YuMin M. Loh, Tai-Ting Lee, Wan-Tze Chen, WenWei Loh, Takuro S. Ohashi, Daniel F. Eberl, Marta Andrés, Matthew P. Su, Azusa Kamikouchi

**Affiliations:** Graduate School of Science, Nagoya University, Nagoya, Japan; Institute of Transformative Bio-Molecules (WPI-ITbM), Nagoya University, Nagoya, Japan; Ear Institute, University College London, 332 Gray’s Inn Road, London, WC1X 8EE, UK; Department of Biology, The University of Iowa, Iowa City, IA, USA; Institute for Advanced Research, Nagoya University, Nagoya, Japan; The Francis Crick Institute, 1 Midland Road, London, NW1 1AT, UK; Animal Health Research Centre, National Institute for Agricultural and Food Research and Technology, Spanish National Research Council (CISA-INIA-CSIC), 28130 Valdeolmos, Spain

**Keywords:** *Aedes* mosquitoes, hearing, neurotransmitters, octopamine, electrophysiology, second messenger

## Abstract

The powerful ears of male *Aedes aegypti* mosquitoes facilitate identification and localization of mating partners *via* detection of female flight tones. Male hearing function is modulated by the efferent release of neurotransmitters, though the secondary mechanisms underlying this modulation remain unclear. Here, we investigated these mechanisms using octopamine as a model, as octopamine modulates hearing function and the erection status of fibrillar hairs lining male ears. We found that pharmacological interference with octopamine receptors alters hearing function at multiple levels and identified the second messenger cAMP as likely mediating these changes. Furthermore, the erection status of male ear fibrillar hairs could be altered by targeting specific sub-types of octopamine receptors, but these changes were not linked to changes in ear frequency tuning. Finally, we suggest that octopamine α2 receptors linked to fibrillar hair erection may not always produce functional proteins across species, with downstream implications for hearing behaviors.

**Highlights:**

- Octopamine and its receptors are found throughout *Aedes aegypti* mosquito ears
- Pharmacological interference with octopamine receptors modulates hearing function
- Modulating levels of the second messenger cAMP also alters hearing function
- Ear frequency tuning does not correlate with the erection of hairs lining male ears

## Introduction

Mosquito-borne diseases represent a major global public health crisis, with over half the world’s population at risk of infection from diseases such as dengue and malaria^1–3^. With disease control measures relying heavily on interventions which reduce mosquito population sizes to inhibit disease transmission, the development of novel control tools with new mechanisms of action/targets are of increasing value.

One such target is mosquito hearing, as males rely on listening for the sounds of flying conspecific females to initiate courtship^4,5^. Male mosquito ears are thus highly sensitive and finely tuned, comprised of a flagellum (sound receiver) whose stiffness can be altered to modulate hearing sensitivity and a cochlea-equivalent known as the Johnston’s Organ [JO]^6,7^. The flagellum is lined with fibrillar hairs, potentially to increase the receiver surface area^8,9^. The JO, which occupies the majority of the second antennal segment (pedicel) in mosquitoes, contains approximately 15,000 sensory neurons in *Aedes aegypti* [*Ae. aegypti*] males^6^, making it the largest in the insect class, as well as an efferent network which allows for transport of neuromodulators from the brain to the ear^10,11^.

Males are able to locate females due to spectral matching between female Wing Beat Frequencies [WBFs] and male flagellar vibrations (ear mechanical tuning)^12^. Intriguingly, male JO neuron responses (ear mechanical tuning) are not particularly sensitive to female WBFs, but instead appeared tuned to distortion products generated from the non-linear mixing of male and female WBFs^12,13^. This mismatch between mechanical and electrical tuning frequencies has been identified in numerous disease-transmitting mosquito species^12–15^.

The auditory efferent network of male mosquitoes has been suggested to modulate hearing sensitivity at both mechanical (flagellar vibration) and electrical (nerve response) levels^11^. For example, altering serotonin levels in *Ae. aegypti* mosquitoes leads to correlated changes in flagellar vibrations, whilst antagonizing octopamine receptors [OctRs] alters both flagellar vibrations and nerve responses^9,10,16,17^. Severing all auditory efferent networks has been shown to induce self-sustained oscillations [SSOs] in male mosquito ears, implicating efferent neurotransmitter release in the modulation of this male specific phenomenon in which male flagellar vibrations increase in displacement by several hundred-fold compared to non-SSO states^18^.

Mosquito ears are actively tuned by JO neurons; However, the anatomical properties of the ear itself can also modulate hearing function. Erection of the fibrillae hairs which extensively cover male flagellae can alter hearing sensitivity in *Anopheles gambiae* [*An. gambiae*] mosquitoes^19^, with increased hearing sensitivity potentially linked to mating behaviors. Male fibrillar hair erection is circadian in *Anopheles* males, with peaks in erection status at dusk (when mating is most likely to occur)^20^, though this has not been described for other species. Investigating differences in fibrillar hair erection status across species is thus crucial in linking hearing systems to mating behaviors.

Prior work in *An. gambiae* has shown that fibrillae states are under the control of octopamine [OA] signalling pathways, with exposure to OctR agonists resulting in rapid erection of these fibrillar hairs^20^. These octopamine signalling pathways can themselves be modulated by GABA signalling pathways which are initiated in the thorax; antagonizing GABA receptors leads to the release of octopamine which binds to OctRs in the flagellum^20,21^. Alpha-like octopamine receptors [OctαRs] have specifically been implicated in these changes in fibrillae erection status, though knockout of a Beta-like octopamine receptor [OctβR] resulted in the permanent collapse of male fibrillae in *An. gambiae*^9^. Interestingly, two orthologs of *Drosophila* Alpha2-like octopamine receptors [Octα2Rs] have been identified in *Ae. aegypti* (AAEL004245 and AAEL025638) whilst only one has been found in *An. gambiae* and *Drosophila melanogaster*^11^. It remains unclear however if this duplication of receptors is related to differences in fibrillae erection status across mosquito species.

The downstream, secondary mechanisms underlying changes in hearing function and fibrillar hair erection following octopamine exposure remain unclear. Given that exposure to the universal second messenger cyclic adenosine monophosphate [cAMP] also leads to fibrillar erection in *An. stephensi* mosquitoes when combined with compounds which inhibit cAMP degradation^20^, changes in fibrillar erection may be linked to the activity of this messenger system following activation of G-Protein Coupled Receptor [GPCR] pathways. However, the exact components of this pathway, as well as fibrillar erection mechanisms in non-*Anopheles* species, remain unclear^9^.

Here, we first screened *Ae. aegypti* octopamine and OctRs expression in three tissues (the pedicel which houses JO, flagellum and the head) *via* immunohistochemistry [IHC] and RT-qPCR. We found that whilst the Beta2-like octopamine receptor [Octβ2R] (AAEL005945) was specifically and abundantly overexpressed in both male and female pedicels compared to the head, Octα2Rs (AAEL004245 and AAEL025638) were instead relatively overexpressed in the flagellum.

Next, we conducted hearing function (mechanical and electrical frequency tuning) assays to identify time course changes arising from octopamine-related compound injection. We found that exposure to the important second messenger cAMP leads to equivalent changes in hearing function resembling octopamine injection, suggesting this messenger type may act within the octopamine signaling pathways to modulate hearing function. In addition to changes in JO neuron function, we also identified changes in male fibrillar erection status following exposure to different octopamine-related compounds. Simultaneous recording of male fibrillar erection status and ear vibrations allowed us to test for potential correlations between fibrillae angle, mechanical tuning and flagellar displacement, with no correlation being identified between these parameters.

Finally, we attempted to link cross-species differences in fibrillae erection status to the presence of one or two orthologous copies of Octα2R in different mosquito species. By conducting cloning and sequencing at genomic DNA and complementary DNA [cDNA] levels in *Aedes aegypti*, we found that pre-mRNA transcripts encoded separately by these two Octα2R genes are spliced into a full-length, mature transcript with predicted protein sequence that shows close alignment with that of encoded by the single copy *An. gambiae* and *D. melanogaster* gene. Given the adjacent position of these two genes on Chromosome I of *Ae. aegypti* (AAEL025638 followed by AAEL004245) and that the two genes independently evolved stop and start codon respectively, we hypothesized that they arose from gene fission/chromosomal rearrangement events that lead to the fragmentation of a single ancestral gene into two. Investigations into other Dipteran species further identified the presence of such splitting events in the Octα2R gene in other mosquito species, with potential implications for their hearing function and behaviors.

## Results

### Octopamine and OctRs are expressed in male and female *Ae*. *aegypti* ears

Mosquito ears are comprised of two components: a JO where auditory mechanotransduction occurs, and a flagellum whose vibrations are actively tuned by JO neurons (Fig 1A; Fig S1A). This flagellum is covered by fibrillar hairs whose erection status has been reported to alter male hearing sensitivity in some mosquito species^19,20^. Previous publications have identified the presence of octopamine release in the pedicels of male *Culex quinquefasciatus*^10^, though its presence and localization in male *Ae. aegypti* pedicels and flagallae remain unclear; octopamine signaling in female mosquitoes have not been reported. We thus first investigated distributions of octopamine throughout *Ae. aegypti* ears in both sexes.

**Figure 1:**
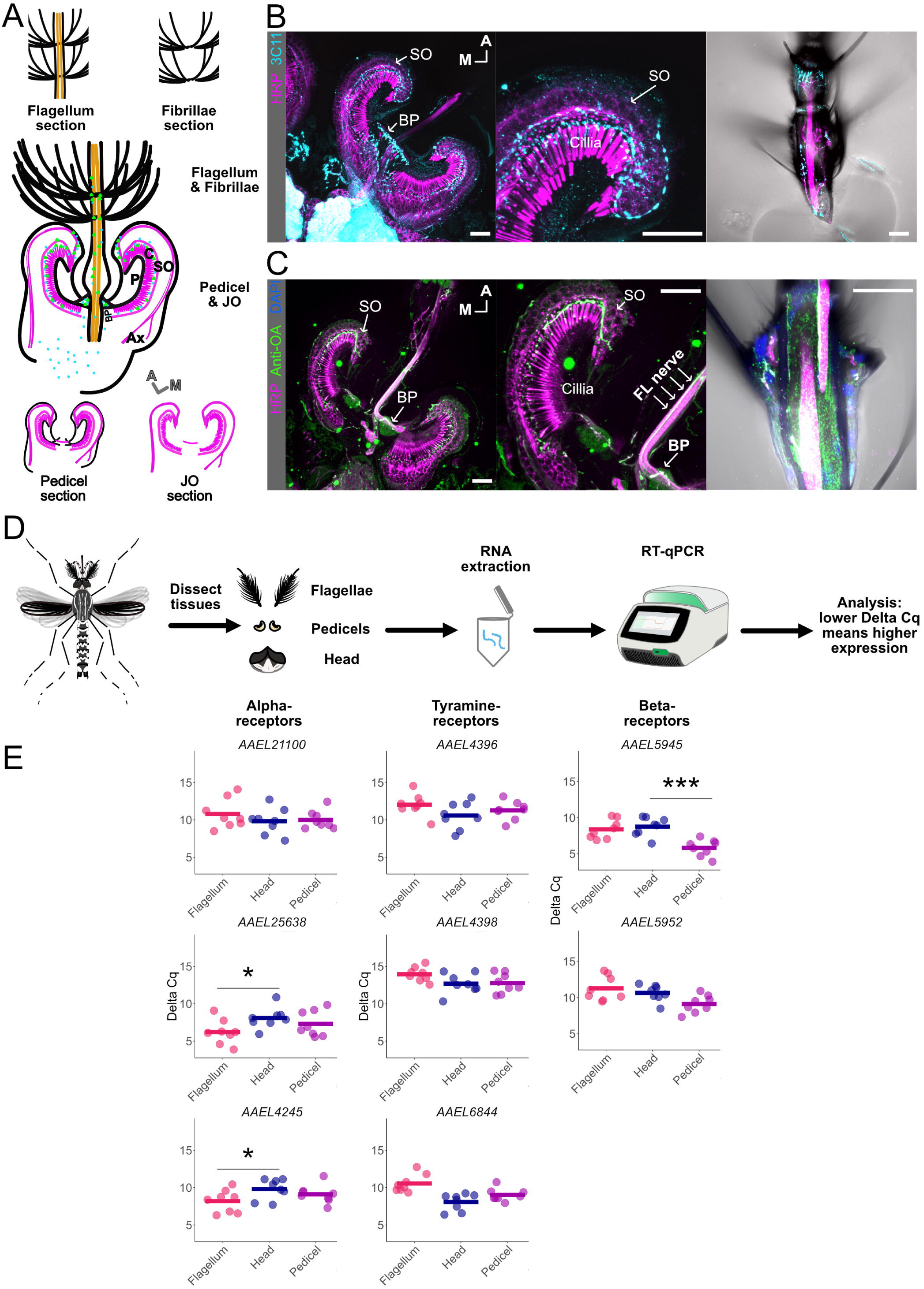
Octopamine potentially overlaps with presynaptic sites and octopamine receptors are expressed in male *Ae. aegypti* flagellar ears. (A) Schematic diagrams of male ear and interpretation of different ear parts: flagellum (top left), fibrillae (top right), pedicel (bottom, left) and JO (bottom right). Magenta and orange represent JO and flagellar neurons in the ear, respectively. Green and cyan dots represent octopamine and presynaptic sites, respectively. Ax, axons of JO neurons; BP, basal plate; C, cilia of JO neurons; P, prong; SO, somata array of JO neurons. A, anterior; M, medial. (B) Location of presynaptic sites in male ear. Sections of male JO (left), JO close-up (middle) and flagellum (right) are shown. DAPI (blue) labels nuclei, Anti-horseradish peroxidase (HRP; magenta) used as neuronal marker, anti-SYNORF1 (3C11; cyan) labels presynaptic sites. Scale bar = 20 μm. (C) Location of octopamine in male ear. Sections of male JO (right), JO close-up (middle) and flagellum (right). Anti-HRP (magenta) used as neuronal marker, anti-octopamine (OA, green) shows octopamine localization. DAPI (blue) was used for flagellum. FL, flagellum. Scale bar = 20 μm. (D) Schematic diagram of RT-qPCR experiments. (E) Delta Cq of each of the eight octopamine-family receptors in male flagellum, head and pedicel. Bar shows median of delta Cq of all repeats. Individual points represent delta Cq of an individual repeat. Lower Delta Cq means a higher expression level. Delta Cq of flagellae (pink), head (blue) and pedicels (purple). *, p ≤ 0.05; ***, p ≤ 0.001; ANOVA on ranks. Eight repeats conducted for each tissue.

We confirmed synaptic terminal distributions in male and female pedicels and flagellae via labeling with a 3C11 (anti-synapsin) antibody which identifies pre-synaptic locations and therefore serves as a marker for the presence of an efferent system within the hearing organ (Fig 1B, Fig S1B&D). JO neuron somata are arrayed in the outer region of the pedicel, from where they extend their cilia radially to the prongs located in the inner side of the pedicel (Fig. 1A, middle). In agreement with previous reports^17,18^, we found sexually dimorphic distributions of 3C11 in the pedicels of male and female mosquitoes, with males having pre-synapses in the outer dendritic region (i.e., basal side of cilium) whilst female 3C11 signal was largely restricted to the somata. In the flagellum, we also found some 3C11 signals in both sexes (Fig 1B, Fig S1B).

Given the sexual dimorphisms in 3C11 expression in male and female pedicels, we next used an anti-octopamine antibody to localize octopamine in male and female pedicels and flagellae. In the pedicel, the octopamine signal was largely confined to the space between the cilia and somata (previously classified as type II efferent terminals^11^) in male JO neurons (Fig 1C, Fig S1E), while signals are enriched in the somata (type IV terminals) of female JO neurons (Fig S1C&E). Only males have apparent octopamine signals in the basal plate region (type V terminals) and flagellar nerve (Fig 1C, Fig S1E).

To confirm if signals were indeed due to octopamine antibody-binding, we imaged a negative control of samples without octopamine antibody application and observed no signal (Fig S1F). Octopamine and pre-synaptic sites marked by 3C11 thus appear to overlap in both male and female JOs (Fig 1 B&C, Fig S1 B-E). In the flagellum, we observed a strong octopamine signal with a highly conserved distribution in the flagellar nerve and base of the fibrillae in both sexes (Fig 1C, Fig S1C).

After confirming octopamine expression patterns in mosquito ears, we next used RT-qPCR to investigate expression patterns of OctRs in different tissues (Fig 1D; primer design provided in Table S1). We found that the Octβ2R (AAEL005945) had significantly greater expression in pedicels compared to heads in both males and females (ANOVA; p < 0.001; Fig 1E; ANOVA; p < 0.05; Fig S2B). Interestingly, different OctRs were overexpressed in male (but not female) flagellae, with the Octα2 family-receptors AAEL004245 (ANOVA; p < 0.05) and AAEL025638 (ANOVA; p = < 0.05) now relatively overexpressed compared to the head (Fig 1E).

Octopamine may not only influence male hearing function but also the WBFs of male and female mosquitoes. We thus also investigated OctRs expression in the mosquito thorax (Fig S2A&B). Whilst all OctRs appear expressed in male and female thoraces, we did not identify overexpression of any specific OctR in male thorax tissues compared to other tissues.

Therefore, whilst octopamine is widely distributed throughout male mosquito ears, pedicels and flagellae show differences in the specific OctRs overexpressed in these tissues compared to the head.

### Injection of OctR agonists and antagonists significantly influences the frequency tuning of male ears

Mosquito hearing function can be functionally characterized at multiple levels, including flagellar vibrations (mechanical tuning) and JO neuron responses (electrical tuning). Although male ear mechanical and electrical tuning ranges partially overlap, males are largely mechanically and electrically sensitive to distinct frequency ranges. Previous work in *An. gambiae* and *Cx. pipiens* has shown that octopamine exposure leads to an increase in male mosquito ear mechanical tuning^9,10^, though the effect on *Ae. aegypti* ears at mechanical and electrical levels remains unknown.

We thus injected compounds that modulate octopaminergic signals to determine the effect on *Ae. aegypti* hearing function by calculating the change in peak mechanical and electrical tuning frequency following injection compared to the pre-injection baseline (Δ Frequency) (Fig 2A&B). The maximum Δ Frequency of each individual mosquito was extracted for quantification. By providing sweep stimulation covering 1-1000 Hz, we estimated the peak frequency tuning of both flagellar vibrations and electrical signaling (Fig 2A; analysis paradigm described in methods and Fig S3A-C).

**Figure 2:**
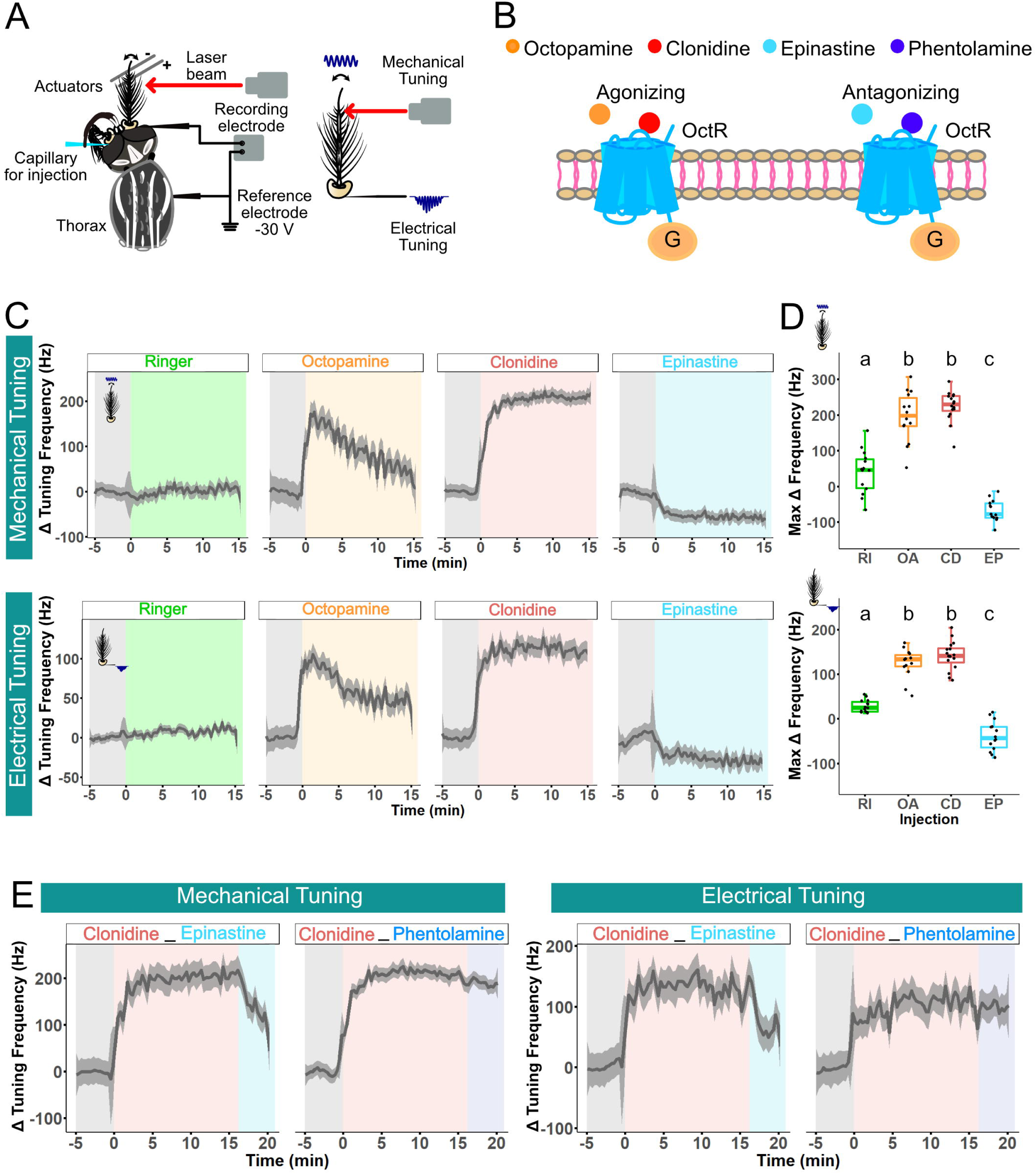
Octopamine alters both mechanical and electrical tuning peak frequencies in male *Aedes aegypti*. (A) Electrophysiology assay diagram. For electrophysiological recording analysis paradigm, see Fig S3A-C. (B) Diagram of mechanism of action for each compound injected. Octopamine (orange), clonidine (red), epinastine (cyan) and phentolamine (blue). (C) Loess fits of mechanical (top) and electrical (bottom) tuning peak frequency changes from each compound injection over 20 minutes. Ringer (green), 1mM octopamine (orange), 1mM clonidine (red) and 1mM Epinastine (cyan). Grey shaded region indicates before injection, colored regions show after injection of compounds. Dark grey shaded area around loess fits indicates 95% confidence interval. Sample sizes: Ringer injection = 14; octopamine injection = 14; clonidine injection= 16; epinastine injection =14. (D) Mechanical (top) and electrical (bottom) max Δ peak frequencies estimates following Ringer, 1mM octopamine, 1mM clonidine and 1mM epinastine injections. Middle line of boxplots represents median values of max mechanical and electrical peak frequency changes from compounds injection. Each dot represents max peak frequency change of each individual mosquito. ns, p > 0.05; *, p ≤ 0.05; **, p ≤ 0.01; ***, p ≤ 0.001; Tukey’s test. Experimental groups denoted by multiple letters are not significantly different from groups containing any of these letters. CD, Clonidine; EP, Epinastine; OA, Octopamine; RI, Ringer. (E) Loess fits of mechanical (left) and electrical (right) tuning peak frequency changes from dual injection over 25 minutes. 1mM clonidine (red) followed by 1mM epinastine (cyan), 1mM clonidine (red) followed by 1mM phentolamine (blue). Grey shaded region indicates before injection, colored regions show after injection of compounds. Dark grey shaded area around loess fits indicates 95% confidence interval. Sample sizes: clonidine followed by phetolamine injection= 4; clonidine followed by epinastine injection =4.

1 mM injection of octopamine into the base of the pedicel caused a rapid increase in mechanical and electrical frequency tuning (Fig 2B&C). A similar increase in both tuning types was observed following injection of the OctR agonist clonidine (CD, 1 mM). Injection of the (potentially beta-like) OctβR antagonist epinastine (EP, 1 mM) however led to a decrease in both mechanical and electrical tuning (Fig 2B&C). The median value of maximum mechanical Δ Frequency through the 20 min after injection was 200 Hz for octopamine, 200 Hz for clonidine and -70 Hz for epinastine, all of which were significantly different from Ringer’s control injections in males (Tukey’s test; p < 0.001; Fig 2D, top; Table 1).

**Table 1:**
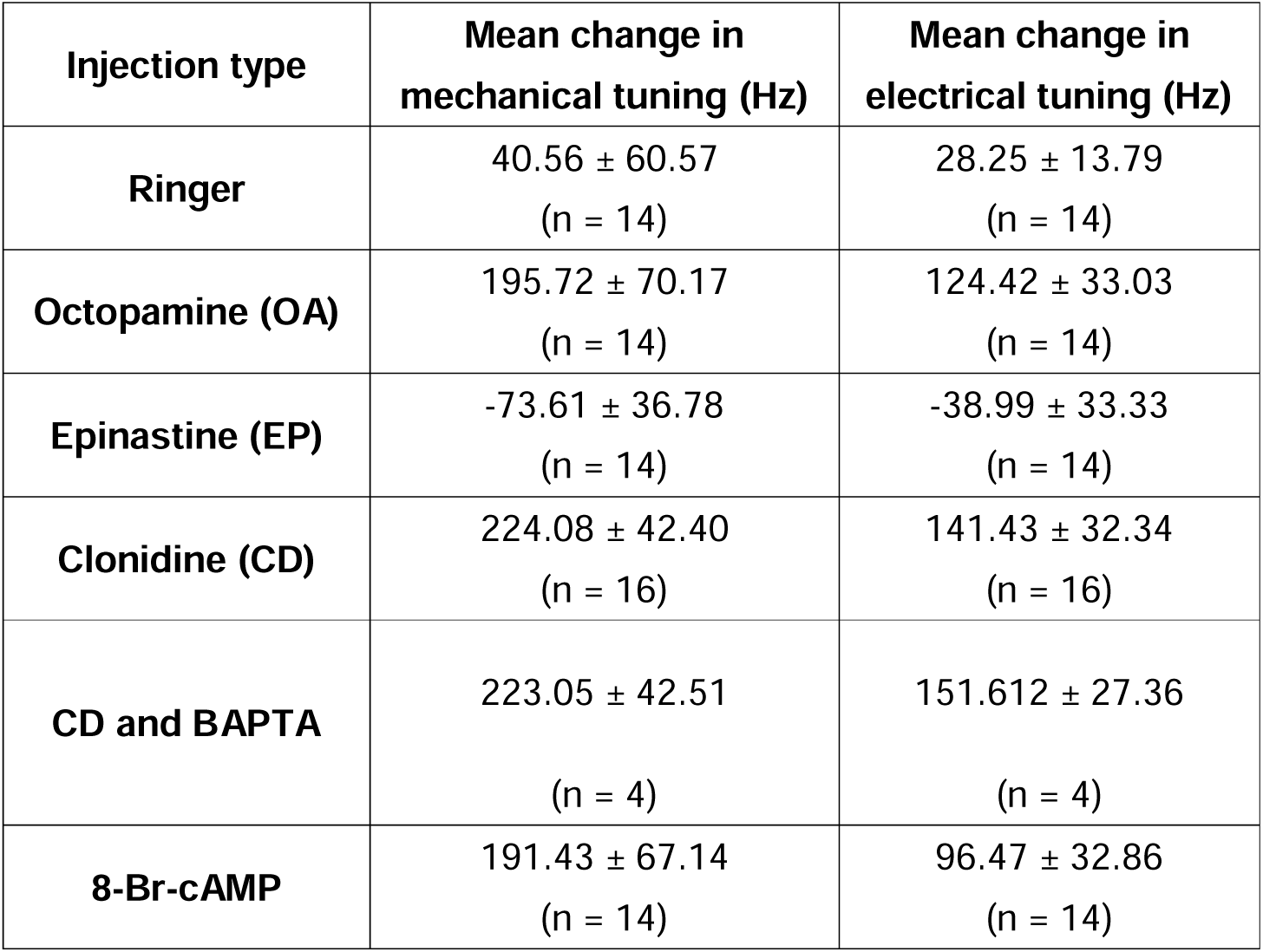
Mean changes in male frequency tuning based on vibrometry/electrophysiology. Mean changes in ear peak mechanical and electrical tuning frequency (based on sweep stimulation analysis) for male *Ae. aegypti* before and after injection of different compounds. Estimated means and standard errors are shown, with sample sizes given in brackets.

Similarly, the median value of maximum electrical Δ Frequency through the 20 min after injection was estimated at ∼ 120 Hz for octopamine, 130 Hz for clonidine and ∼ -50 Hz for epinastine as compared with the Ringer’s control group (Tukey’s test; p < 0.001; Fig 2D, bottom; Table 1). Injection of epinastine about 15 min after clonidine exposure led to a substantial decrease in mechanical and electrical Δ Frequency; however, injection of the (potentially alpha-like) OctαR antagonist phentolamine after clonidine did not result in such a decrease (Fig 2E).

1mM injection of octopamine in females also showed statistically significant differences compared to Ringer control, though these changes were far smaller than the differences calculated for males (Fig S3D&E).

Injection of octopamine or OctR agonists thus leads to a reproducible increase in peak mechanical and electrical tuning in both sexes, whilst injection of specific OctR antagonist in males results in a decrease in peak frequency tuning (examples shown in Fig S4A&B).

### Modulating the level of the second messenger cAMP influences male *Aedes aegypti* mechanical and electrical frequency tuning

Binding of octopamine to GPCRs leads to altered regulation of the second messenger cyclic adenosine 3′,5′-monophosphate (cAMP), as well as other second messenger systems (Fig 3A; ^22,23^). Prior work has shown that direct exposure of 8-Br-cAMP (a cell-permeable analog of cAMP), in addition to theophylline to prevent cAMP degradation, alters male *An*. *stephensi* fibrillar hair erection status^20^, suggesting a role for this messenger in regulating mosquito hearing sensitivity; its role in mediating frequency tuning remains unclear.

**Figure 3:**
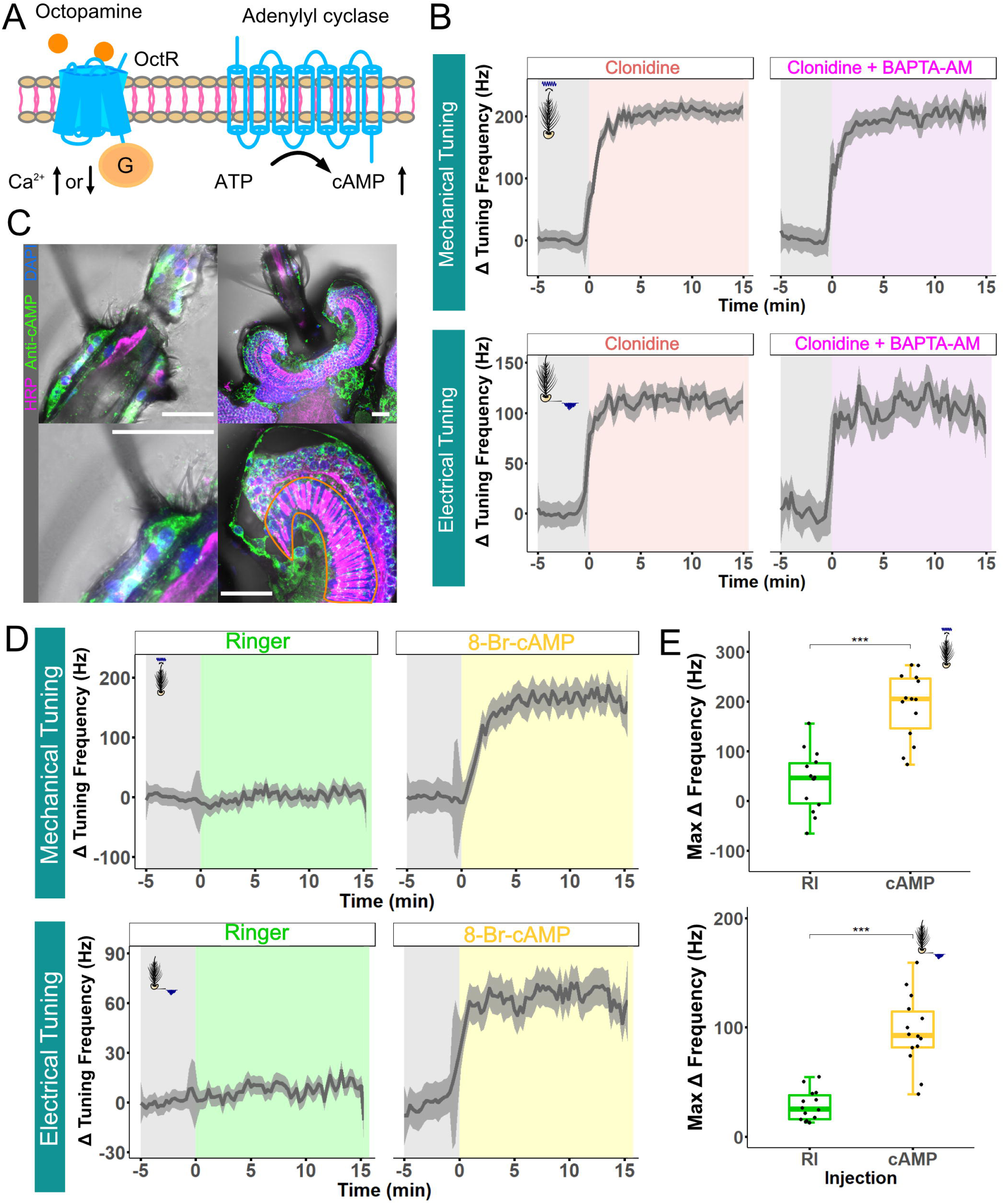
Octopamine potentially modulates male mosquito fibrillae status and mechanical frequency tuning via different second messengers. (A) Diagram of second messenger pathways activated inside cells following binding of octopamine to relevant GPCRs. (B) Loess fits of mechanical (top) and electrical (bottom) tuning peak frequency changes of male mosquitoes. 1mM Clonidine alone (red), 1mM clonidine plus BATPA-AM (purple). Grey shaded region indicates before injection, colored regions show after injection of compounds. Dark grey shaded area around loess fits indicates 95% confidence interval. Sample sizes: Clonidine injection= 16; Clonidine+BAPTA-AM injection = 4. (C) cAMP signal in male ear, male flagellum (top left), flagellum close-up (bottom left), JO (top right) and JO close-up (bottom right). HRP staining (magenta) allows for visualizing neurons, DAPI labels nuclei (blue) and anti-cAMP signal (green) shows cAMP distribution in male ear. Scale bar = 20 μm. Orange line indicates region containing ciliated sites. (D) Loess fits of mechanical (top) and electrical (bottom) tuning peak frequency changes of 8-Br-cAMP injection and Ringer control over 20 minutes. Ringer (green) and 1mM 8-Br-cAMP (yellow). Grey shaded region indicates before injection, colored regions show after injection of compounds. Dark grey shaded area around loess fits indicates 95% confidence interval. Sample sizes: Ringer injection = 14; 8-Br-cAMP injection = 14. (E) Mechanical (top) and electrical (bottom) max Δ peak frequencies change of Ringer and 8-Br-cAMP injections. Middle line of boxplots represents median values of max mechanical and electrical peak frequency changes from compounds injection. Each dot represents max peak frequency change of each individual mosquito. ns, p > 0.05; *, p ≤ 0.05; **, p ≤ 0.01; ***, p ≤ 0.001; Tukey’s test.

First, we injected male mosquitoes with 1mM clonidine mixed with the cell-permeant Ca^2+^ chelator-BAPTA-AM and found no differences in changes to mechanical and electrical frequency tuning compared to clonidine injection alone (Fig 3B). This suggests that Ca^2+^ may not be responsible for the previously observed changes in male mosquito hearing tuning resulting from octopamine injection.

We next used IHC to localize cAMP in mosquito ears and found abundant cAMP signals in the somata of male JOs and flagellae, which were aggregated around nuclei (Fig 3C). Limited cAMP signal was also found in the region which co-localizes with ciliated dendrites (Fig 3C, region highlighted with orange line).

Injecting 1mM 8-Br-cAMP into male mosquitoes induced similar changes in mechanical and electrical tuning as octopamine (Fig 3D), with mechanical and electrical median value of max Δ Frequency ∼ + 190 Hz and ∼ + 95 Hz (Tukey’s test, p < 0.001, Fig 3E). cAMP thus appears to play a major role in modulating changes in ear mechanical and electrical tuning following octopamine exposure.

### Injection of OctR agonists and antagonists alters the fibrillae erection status of male flagellar ears

Male antennal fibrillae erection status has been linked to hearing sensitivity modulation in *An. gambiae* mosquitoes^9,19^. A previous study found a unique structure (referred to as an annulus) above the fibrillae base in male *Anopheles* mosquitoes which seemingly induced fibrillae erection status via conformational changes^24^. Although *Ae. aegypti* fibrillae have been generally regarded as erect at all times, our male *Ae. aegypti* flagellae IHC data also found this annulus in two conformational states depending on the fibrillae erection status (Fig S5A), suggesting that male *Ae. aegypti* fibrillae status is potentially changeable more flexible than previously assumed and may also show changes across the day (as previously reported for *Anopheles* species). We did not observe equivalent structures in female flagellae (Fig S5A). In our experiments, preparation of mosquitoes for electrophysiology experiments can result in collapsed fibrillae in males, enabling injection of compounds to investigate changes in fibrillae erection status.

To investigate if *Ae. aegypti* fibrillae status is correlated with mechanical frequency tuning and displacement, we recorded fibrillae movement and unstimulated mechanical tuning frequency simultaneously before and after compound injection (Fig 4A). Angular changes in fibrillae movement were then quantified using a DeepLabCut [DLC] based methodology (Fig S5B). The unstimulated flagellar frequency change (Δ Unsti Frequency) and angle change (Δ Angle) was calculated by post injection unstimulated frequency and angle minus the pre-injection median baseline, respectively. Then, maximum unstimulated frequency change (Max Δ Unsti Frequency) and angle change (Max Δ Angle) were extracted for quantification.

**Figure 4:**
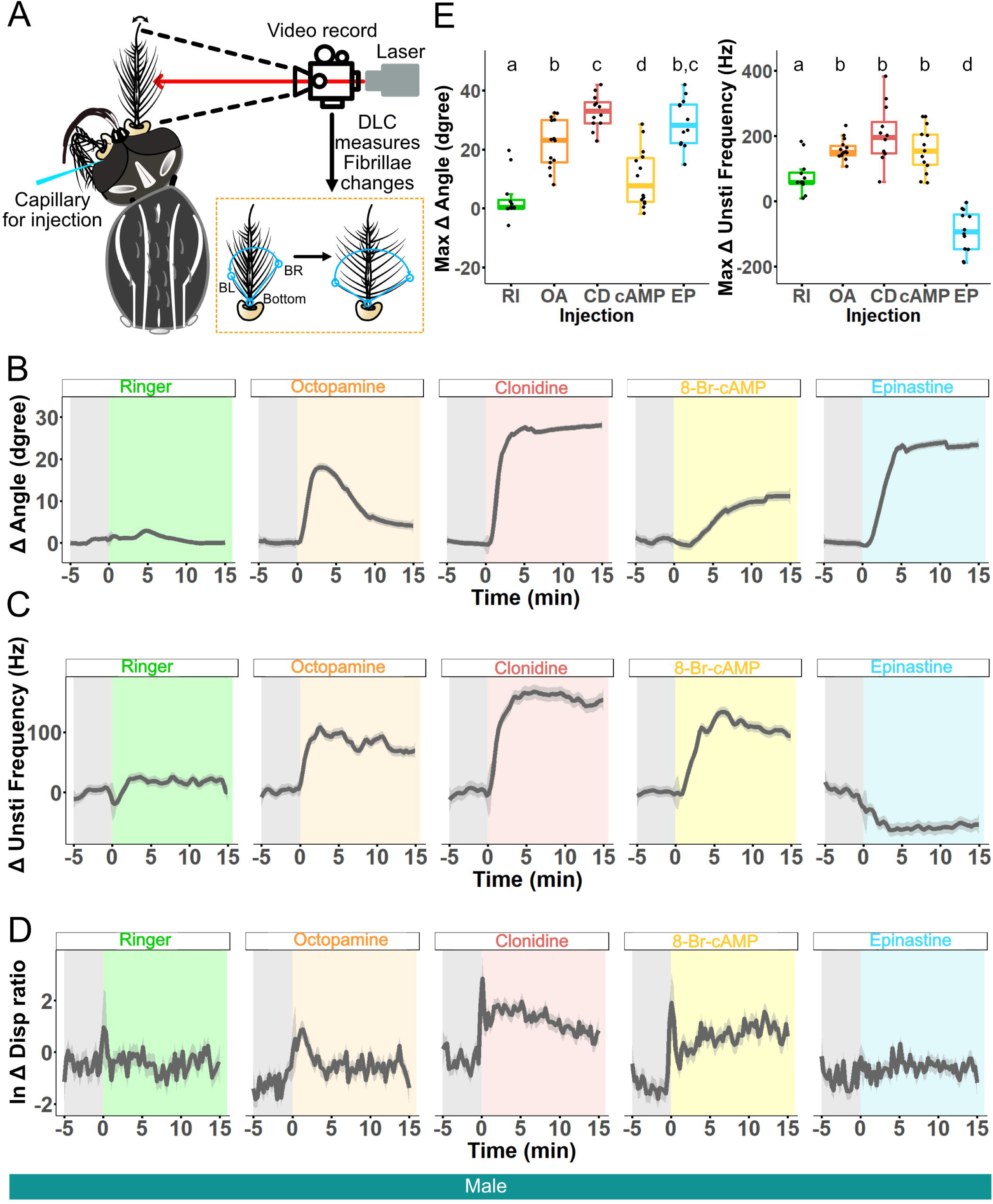
Octopamine modulates male *Ae. aegypti* fibrillae status and peak mechanical frequency tuning. (A) Schematic of DLC experimental analysis. Left: Unstimulated mechanical tuning frequency and fibrillae status are recorded by laser Doppler vibrometry and camera build in laser, respectively. (B) Loess fits of changes in fibrillae angle relative to flagellum following compound injection over 20 minutes. Ringer (green), 1mM octopamine (orange), 1mM clonidine (red), 1mM 8-Br-cAMP (yellow) and 1mM epinastine (cyan). Grey shaded region indicates before injection, colored regions indicate after injection of compounds. Dark grey shaded area around loess fits indicates 95% confidence interval. Sample sizes: Ringer injection = 12; octopamine injection = 14; clonidine injection= 12; 8-Br-cAMP = 14; epinastine injection = 12. (C) Loess fits of change in unstimulated mechanical tuning frequency following compound injection over 20 minutes. Ringer (green), 1mM octopamine (orange), 1mM clonidine (red), 1mM 8-Br-cAMP (yellow) and 1mM epinastine (cyan). Grey shaded region indicates before injection, colored regions indicate after injection of compounds. Dark grey shaded area around loess fits indicates 95% confidence interval. Sample sizes: Ringer injection = 12; octopamine injection = 14; clonidine injection= 12; 8-Br-cAMP = 14; epinastine injection = 12. (D) Loess fits of log ratio change in unstimulated flagellar displacement following compound injection over 20 minutes. Ringer (green), 1mM octopamine (orange), 1mM clonidine (red), 1mM 8-Br-cAMP (yellow) and 1mM epinastine (cyan). Grey shaded region indicates before injection, colored regions indicate after injection of compounds. Dark grey shaded area around loess fits indicates 95% confidence interval. Sample sizes: Ringer injection = 12; octopamine injection = 14; clonidine injection= 12; 8-Br-cAMP = 14; epinastine injection = 12. (E) Quantification of changes of max fibrillae angular movement (left) and max unstimulated mechanical tuning frequency change (right) following compound injection. Middle line of boxplots represents median value, whilst each dot represents data from an individual mosquito. ns, p > 0.05; *, p ≤ 0.05; **, p ≤ 0.01; ***, p ≤ 0.001; Pairwise wilcox test. Experimental groups denoted by multiple letters are not significantly different from groups containing any of these letters.

We found no change in fibrillae status after Ringer injection (Fig 4B; Table 2). Both 1mM octopamine and clonidine caused male fibrillae to become more fully erect compared with Ringer controls (Pairwise Wilcox test, p < 0.001, Fig 4B&E). However, 1mM epinastine injection also resulted in significantly greater erection of male flagellae despite significantly reducing the unstimulated mechanical frequency tuning (Pairwise Wilcox test, p < 0.001, Fig 4C&E; Table 2).

**Table 2:** Median changes in unstimulated male peak mechanical frequency tuning and fibrillae erection angle. Median changes in ear peak mechanical frequency as well as median fibrillae angle (based on analysis of unstimulated recordings) for male *Ae. aegypti* before and after injection of different compounds. Estimated medians and standard errors are shown, with sample sizes given in brackets.

| Injection type | Median change in unstimulated mechanical tuning (Hz) | Median change in fibrillae angle (degree) |
| --- | --- | --- |
| Ringer | 58.67 ± 15.38<br>(n = 12) | 0.69 ± 2.03<br>(n = 12) |
| Octopamine (OA) | 153.29 ± 7.96<br>(n = 14) | 23.07 ± 2.23<br>(n = 14) |
| Epinastine (EP) | -97.08 ± 18.51<br>(n = 12) | 28.31 ± 2.41<br>(n = 12) |
| Clonidine (CD) | 201.22 ± 24.60<br>(n = 12) | 32.96 ± 1.66<br>(n = 12) |
| 8-Br-cAMP | 154.61 ± 18.35<br>(n = 14) | 7.74 ± 2.63<br>(n = 14) |

Interestingly, 1mM 8-Br-cAMP injection resulted in relatively small (though still significant) changes in male fibrillae erection status compared to Ringer (Pairwise Wilcox test, p < 0.05, Fig 4B&E) despite the significant increase in mechanical tuning observed (Pairwise Wilcox test, p < 0.001, Fig 4C&E).

Injection of 1mM octopamine into female mosquitoes resulted in no significant change in fibrillae status compared to Ringer injected controls (Fig S5F top, ns, p > 0.05, t test), though the unstimulated mechanical frequency tuning did significantly increase (Fig S5F bottom, t test).

We next calculated the effect of compound injection of flagellar displacement (Δ Disp) via post injection displacement minus pre injection baseline, and found that octopamine, clonidine and 8-Br-cAMP injections showed immediate, sustained, increases in flagellar displacement (Fig 4D). No obvious change in flagellar displacement was observed following epinastine injection (Fig 4D).

Finally, we calculated the correlation between fibrillae angle and unstimulated mechanical tuning frequency before and after injection (Fig S6A). Before injection, there were no strong correlation between the two values; whilst agonists/second messengers tended to increase the correlation after injection while antagonists decreased it, only clonidine injection resulted in a statistically significant difference in correlation between pre and post injection status (Fig S6A, p < 0.05, Wilcoxon Signed-Rank Test). In general, we found no correlation between fibrillae angular status and unstimulated mechanical tuning frequency. Furthermore, we found no correlation between fibrillae status and flagellar displacement in all injection groups (Fig S6B, ns, p > 0.05, Pared t test for males, Wilcoxon Signed-Rank Test for females).

### *Ae. aegypti* Octα2Rs are the result of splitting of single ancestral receptor

OctαRs have previously been linked to changes in mosquito hearing function and ear fibrillar erection^9,20^. Our RT-qPCR data demonstrates that among all OctRs, both Octα2Rs (AAEL025638 and AAEL004245) showed the highest expression in the male flagellum. The presence of two orthologous Octα2Rs in the genome as compared to the single copy identified in *An. gambiae* and *Drosophila melanogaster* [*D. melanogaster*]^11^ is also distinct in *Ae. aegypti*. Given that male *Ae. aegypti* fibrillae have generally been regarded as permanently erect, as compared to *An. gambiae* males which show timely-dependent fibrillae erection/collapse, we wondered if the gene copy numbers of Octα2Rs could be linked to species-specific differences in fibrillae erection physiology.

^11^Using the Liverpool L5 genome of *Ae. aegypti*, we found that not only were the coding sequences for the two Octα2Rs located sequentially on the complementary strand of chromosome 1 (Fig S7A), but that each of these genes contained a reduced number of predicted transmembrane domains [TMDs] essential for GPCR function; In contrast to the seven TMDs found for canonical GPCRs, DEEPTMHMM^25^ analysis on the predicted protein sequences of the two Octα2Rs found that AAEL025638 only contained four TMDs and AAEL004245 three TMDs (Fig S7B). Protein sequence alignment with the predicted protein sequences of *D. melanogaster* and *An. gambiae* single copies of Octα2R further found that the four TMDs of AAEL025638 aligned to the first half of the predicted full-length sequences of the latter two species while the three TMDs of AAEL004245 aligned to the second half of the sequences, suggestive of a single origin for the two *Ae. aegypti* Octα2Rs.

Closer scrutinization found that AAEL025638 and AAEL004245 have independently evolved stop and start codons, respectively (Fig S7C). These observations therefore suggest that if the two Octα2Rs were transcribed and translated independently, neither would form a functional seven transmembrane GPCR, unlike that of *An. gambiae* and *D. melanogaster*, supported by GPCRHMM predictions.

Given that AAEL025638 and AAEL004245 are sequentially located on Chromosome 1, it is possible that these genes have been incorrectly annotated and are instead a single gene with an unusually long intron (∼775,000 base pairs). To test this, we designed primers for RT-PCR and sequencing around the end of AAEL025638 and start of AAEL004245. We unexpectedly found that we were able to obtain a full-length cDNA clone (∼1.8kbp long) closely resembling the transcript length of the single copy Octα2R gene in *An. gambiae* and *D. melanogaster* by using a forward primer that binds to the start of the coding sequence (CDS) of AAEL025638 and the end of the CDS of AAEL004245. Sanger-sequencing of the cDNA clone revealed that the cDNA sequence is a joint-sequence that contains two sequence fragments that separately mapped to the CDS of AAEL025638 and AAEL004245 (Fig S7D & E).

Whilst neither of the predicted protein sequences of the two genes were considered GPCRs based on GPCRHMM predictions^26^, the translated protein sequence of the full-length cDNA was now considered a GPCR (Fig S7F) and showed close alignment with the Octα2R protein sequences of *An. gambiae* and *D. melanogaster*. Interestingly, we found that the last exon of AAEL025638 and first exon of AAEL004245 were spliced out from the full-length cDNA sequence, suggesting that a *trans-splicing* event could be underlying the splicing and joining of two independent pre-mRNAs into a full-length mature mRNA. We also cloned out and sequenced PCR fragments corresponding to different regions of AAEL025638 and AAEL004245 from genomic DNA to confirm the accuracy of the genome. However, we could not exclude the possibility that these exons have been incorrectly annotated, and that instead AAEL025638 and AAEL004245 exist as a single gene.

We next tested if two copies of Octα2R had also been identified in other Dipteran species. We used protein BLAST to identify the 5000 closest orthologs of *D. melanogaster*’s Octα2R and tested if these hits were categorized as GPCRs based on GPCRHMM (Supplemental Files 1 & 2). We then used MMseqs2^27^ to cluster these protein sequences of these 5000 hits and isolated four clusters (*D. melanogaster* Octα2R cluster, *An. gambiae* Octα2R cluster, as well as AAEL025638 and AAEL004245 clusters (Fig S8A)). Analyses of the AAEL025638 and AAEL004245 clusters identified nine mosquito species in which Octα2R had seemingly been split in two (Fig S8B; Table S3), though it is possible that all species may have single copies of Octα2R with long introns spanning hundreds of thousands of base pairs. In general, *Anopheles* species tended to have single copies of Octα2R (although *Anopheles coustani* and *Anopheles marshalli* have two), whilst species from the tribe Aedini (including *Aedes aegypti*, *Aedes albopictus*, *Armigeres subalbatus* and *Ochlerotatus camptorhynchus*) all had two copies (Fig S8C).

We attempted to reconstruct full-length Octα2R sequence from the translated CDS of the two genes for these nine mosquito species using *D. melanogaster* and *An. gambiae* protein sequences as templates and observed considerable variation around the stop and start codons across species (Fig S8C & D), suggesting divergent evolution of these translational termination and start sites across these mosquito species. Interestingly, the start of the *Aedes albopictus* ortholog of AAEL004245 lacks a crucial sequence of amino acids which code for proper GPCR function (Fig S8E). As such, even if mRNA from the two sequential genes can be spliced together, a functional version of Octα2R protein may not be produced.

By looking at the gene structure of Octα2R across species, we found that the two CDS fragments for species with split Octα2R when aligned with those species with one Octα2R, consistently mapped to two adjacent exons with no overlapping. We therefore investigated the intron bridging these two specific exons for species with one Octα2R within the *D. melanogaster* Octα2R cluster and *An. gambiae* Octα2R cluster: We found members of the *An. gambiae* Octα2R cluster displayed longer intron length compared to members of the *D. melanogaster* Octα2R cluster, suggesting intron expansion from Drosophilidae to Anopheline, which could be an indication of chromosomal rearrangement events occurring within this intronic region that eventually lead to gene fragmentation for other mosquito species (Supplemental File 1).

To link these molecular results with behavioral observations of male fibrillae erection status, we searched the literature for all previous reports on mosquito fibrillae erection status. We identified publications covering 23 different mosquito species categorizing mosquitoes into three groups: ‘Permanently erect’, ‘Erect/Collapsed’ and ‘Short hairs’ (Table S4). Whilst many reported species did not have associated genomic data, we note that for those species with reported genomic data, splitting of Octα2R was often linked to relatively permanent fibrillae erection.

## Discussion

Hearing represents a promising target for novel methods of mosquito control given the important role it plays during mating across many disease-transmitting species^28^. Recent molecular work has built atop decades of anatomical and physiological reports to substantially improve our understanding of the fundamental mechanisms underlying male hearing function in several species^9,12,18^. However, given the significant diversity observed in mosquito hearing systems across species, as well as the multiple interlocking components underlying hearing function, further species-specific research is needed.

Previous profiling of serotonin distributions in male and female ears did not identify sexual dimorphisms in their expression patterns within the JO, despite differences in 3C11 localization; in both sexes, serotonin was expressed only in the somata of JO neurons^17^. Here however we found that octopamine signals were clearly distinct between the sexes (Fig 1C, Fig S1C&E), with male octopamine expression potentially overlapping with regions containing 3C11 (Fig S1D&E), providing further support to the efferent nature of neurotransmitter release within the mosquito JO. The abundance of octopamine identified in the flagellae of males and females (Fig 1C, Fig S1C) could be linked to olfactory processes (as in other insects^29^), as well as fibrillae erection in males.

This erection has been linked in *An. stephensi* to changes in morphology of an annulus located directly above each fibrillae^24^. Our flagellar sections clearly identified equivalent structures in males but could not see such annuli in females (Fig S5A). Sexual dimorphisms in the developmental bases of such structures, as well as differences across species, could therefore be investigated in the future. Furthermore, the location of OctRs in these structures requires further research. Circadian release of octopamine has been hypothesized to underlie the circadian nature of male fibrillae erection^9^ and may thus alter hearing function. The circadian mechanisms underlying octopamine release could also represent a potential target to interfere with mosquito mating.

Previous RNAseq analyses in *An. gambiae* identified Octβ2R as having the highest expression of all OctRs in male pedicels^9^. Our qPCR data suggests this is also the case in *Ae. aegypti* pedicels, with AAEL005945 significantly overexpressed in male pedicels compared to heads (Fig 1E). AAEL005945 was not overexpressed in the flagellum compared to the head in males however, with the two *Ae. aegypti* Octα2R orthologs instead significantly higher in expression compared to heads (Fig 1E). Prior work in *An. stephensi* has linked alpha-type receptors to fibrillae erection^20^, suggesting differential roles for OctRs in different tissues (though disruption of Octβ2R did result in permanent loss of fibrillae erection in *An. gambiae*^9^).

Female mosquitoes exposed to high (80 mM) concentrations of clonidine showed flight deficiencies compared to controls, suggesting some role for octopamine in mosquito flight^30^. We however did not identify overexpression of any OctRs in the thorax compared to other tissues (flagellum, head, and pedicel) in both sexes (Fig S2). Previous work in *Drosophila* has found significant differences in flight initiation and maintenance in flies lacking the octopamine synthesis enzyme tyramine-β-hydroxylase, though differences in WBF were not observed^31^. Investigation into more specific tissue types (such as indirect flight muscle) is necessary to test the significance of octopamine for mosquito flight modulation.

Injection of octopamine, as well as the OctR agonist clonidine, led to significant increases in both peak mechanical and electrical tuning (Fig 2C; Table 1). These changes could be potentially mediated *via* Octβ2R, particularly as injection of the supposedly beta-specific antagonist reduced both frequency tuning types whilst injection of the supposedly alpha-specific antagonist phentolamine did not (Fig 2E)^23^. However, the specificity of these compounds when injected at such concentrations is unclear, and off-targets may play confounding factors. Indeed, the differential effects of alpha- and beta-like receptors on intracellular cAMP and Ca^2+^ levels remain unclear in mosquitoes and require the use of cell-based assays for further clarification^32,33^.

The mosquito pedicel houses thousands of JO neurons, with different JO neuronal populations have different electrical frequency tuning^34–37^. Here, we found that octopamine (or OctR agonist) injection resulted in the silencing of lower frequency neurons and thus an overall increase in the median peak electrical tuning frequency (Fig 2D; Fig S4), matching prior reports^16^. This may be due in part to changes in the peak mechanical tuning frequency – if flagellar vibrations are also reduced at lower frequencies (Fig S4), this will likely directly influence electrical tuning frequency range given the coupling between the two tuning types^38^. It remains unclear however how these neuronal populations are distributed in the mosquito JO, and the related expression patterns of OctRs.

8-Br-cAMP injection significantly altered male hearing function (Fig 3D&E, Fig S4). cAMP is a universal second messenger with many different potential downstream signalling pathways, making further elucidation of the underlying mechanisms challenging^39,40^. Indeed, given the universal nature of cAMP, it is likely other neurotransmitters may alter hearing function via equivalent pathways (i.e. serotonin binding to serotonin receptors should also alter cAMP levels)^41,42^. One potential pathway is via binding to cAMP-sensitive channels located within the cell that can modulate ionic concentrations^43^. Alternative pathways have been reported for other types of microtubule bending however, with bending of sperm flagellae linked to cAMP-mediated activation of specific GTPases^44^.

cAMP has been reported to modulate fibrillae erection in male *An. stephensi^20^*. We observed slight modulation for *Ae. aegypti* males (Fig 4B), though to a far lesser degree than octopamine itself. Whilst there may be differences in the underlying pathways, we note that cAMP was only able to result in substantial fibrillar erection in the previous study when coupled with theophylline^20^, which itself can activate a variety of distinct pathways separate from its role in inhibiting cAMP degradation. Given that fibrillar erection appears to be the result of changes in ionic concentration^24^, other factors may instead be potentially relevant.

Whilst injection of OctR agonists and antagonists clearly resulted in differential effects on mechanical and electrical tuning (Fig 2C, Fig 4C), their effect on fibrillae erection appeared conserved (Fig 4B, Table 2); thus, erection status appears not to be correlated with ear frequency tuning following manipulation of the octopamine signaling pathway in male *Ae. aegypti* ears (Fig 4B&C, Fig S6A). This apparent uncoupling of mechanical frequency tuning to fibrillae status was also observed for flagellar displacement, with the substantial increases in flagellar displacement observed following injection of octopamine, clonidine and cAMP apparently not correlated with fibrillae angle change (Fig 4B&D, Fig S6B).

A recent single nucleus transcriptomic atlas of *Ae*. *aegypti* mosquitoes enables visualization of the expression levels of Octα2R (AAEL025638) and Octβ2R (AAEL005945) in different cell types across tissues^45^. Though mosquito pedicels were not included as a distinct tissue type, by filtering by *inactive* (AAEL020482, *iav*) positive cells in the mechanosensory cell type of the head tissue type (head with antennae, proboscis and brain) we isolated putative JO neurons (Fig S9A&B, Fig S10A&B). These putative JO neurons contain significant levels of Octβ2R expression in both sexes but limited Octα2R co-expression (Fig S9C&D, Fig S10C&D), supporting our electrophysiology data which shows that agonizing/antagonizing OctβR leads to a significant change in ear frequency tuning. These findings together could suggest Octβ2R dominates frequency tuning in *Ae. aegypti* JOs.

Moreover, according to the flagellum cell maps, both Octα2R and Octβ2R are overwhelmingly expressed in male cells marked by *ppk317*^46–48^ (Fig S11), a subfamily of the *pickpocket* (PPK) ion channel (DEG/ENaC) family which localizes close to the fibrillae and is expressed in males only^45^. This supports our DLC data, which indicates that Octα2Rs and Octβ2R may both modulate changes in fibrillae state. Further testing of genes overexpressed in this specific cell group could identify further targets involved in fibrillae state changes.

Our displacement analyses found that octopamine, clonidine and cAMP exposure can induce large flagellar displacement (Fig 4D, Table 2). Previous studies categorised the mono-frequent large flagellar displacements as SSOs^9^ but did not correlate time-dependent changes in flagellar displacement to hearing function directly. Our analyses assessing correlations between different hearing function properties (i.e., frequency, displacement, angle) suggest limited correlation between all parameters tested (Fig S6).

Our analyses of Octα2R across Dipteran species found multiple mosquito species in which Octα2R has seemingly been split in two (Fig S8B, C). Our *Ae. aegypti* sequencing data suggests that rather than being a sequencing artefact, this split could be the result of chromosomal rearrangements (Fig S7), though we cannot rule out that these genes are instead a single gene with a long intron. *trans*-splicing of pre-mRNA has been demonstrated for a number of *Drosophila* genes, though further research is needed in mosquito species to test if this mechanism underlies our sequencing results^49^. Given the variation we observed around the start and stop codons of the first gene and second gene respectively across species with two Octα2R orthologs, particularly for *Aedes albopictus*, it appears there is significant selection pressure exerted on these regions, indicating an important role for Octα2R in determining mosquito fitness (Fig S8B-D).

Though the molecular mechanisms underlying SSOs remain unclear, severing of auditory efferent pathways *via* injection of tetanus toxin has previously been reported to also induce SSO onset, suggesting these efferent networks in some way modulate SSOs^18^. Octopamine modulates hearing function across multiple species of disease-transmitting mosquitoes^9,10,16^. The widespread availability of pharmacological agonists/antagonists targeting OctRs^9,10,16,50^, combined with octopamine’s high potency, makes this a potential route for future research into possible mechanisms of mosquito control.

### Limitations of the study

The use of compounds for injection raises the risk of off-target effects. Whilst we attempted to mitigate these risks by testing a variety of both agonists and antagonists, we cannot exclude the possibility of functional changes being the result of binding to non-OctRs. Testing of mutant mosquito lines is necessary to validate our results. Whilst cAMP appears heavily involved in the molecular pathways underlying changes in hearing function after octopamine injection, the multitude of potential second pathways involving cAMP renders it challenging to fully elucidate which pathway(s) are relevant for ear frequency tuning. Finally, our computational analyses of Octα2R are insufficient to directly demonstrate a role for this receptor type in modulating differences in ear fibrillae state across species. They are also limited by a lack of genomic information for many species which have previously been reported to show changes in fibrillae erection status, whilst available genomes may also have issues resulting from insufficient sequencing depth or the production of *de novo* genomes from RNAsequencing data. Indeed, the apparent splitting of Octα2R in 10 Dipteran species may be an artefact of incorrect annotations rather than a real splitting event. Further, cross-species experiments are necessary to validate these analyses.

## Abbreviations

A: anterior
*A. aegypti*: *Aedes aegypti*
*A. albopictus*: *Aedes albopictus*
*An. arabiensis*: *Anopheles arabiensis*
*An. coluzzii*: *Anopheles coluzzii*
*An. coustani*: *Anopheles coustani*
*An. darlingi*: *Anopheles darlingi*
*An. funestus*: *Anopheles funestus*
*An. gambiae*: *Anopheles gambiae*
*An. machlipalpis*: *Anopheles machlipalpis*
*An. marshallii*: *Anopheles marshallii*
*An. merus*: *Anopheles merus*
*An. moucheti*: *Anopheles moucheti*
*An. nili*: *Anopheles nili*
*An. stephensi*: *Anopheles stephensi*
*ASubRV.*: *Armigeres subalbatus*
Ax: axons of JO neurons
BP: basal plate
C: cilia of JO neurons
cAMP: cyclic adenosine monophosphate
CD: Clonidine
cDNA: complementary DNA
CDS: coding sequence
*Cx. p. pallens*: *Culex pipiens pallens*
*Cx. quinquefasciatus*: *Culex. quinquefasciatus*
DLC: Deeplabcut
*D. melanogaster*: *Drosophila melanogaster*
EP: Epinastine
FL: flagellum
GPCR: G-Protein Coupled Receptor
HRP: horseradish peroxidase
*iav*: *inactive;*
IHC: immunohistochemistry
JO: Johnston’s organ
LD: Light:Dark
M: medial
*Mal. genurostris*: *Malaya genurostris*
NGS: normal goat serum
OA: Octopamine
*Oc. camptorhynchus*: *Ochlerotatus camptorhynchus*
OctRs: Octopamine receptors
OctαRs: Alpha-like octopamine receptors
OctβRs: beta-like octopamine receptor
P: prong
PBS: Phosphate-buffered saline
PBT: Phosphate-buffered saline with Triton X-100
PFA: paraformaldehyde
PL: Phentolamine
*ppk*: *pickpocket*
RI: Ringer
*Sa. cyaneus*: *Sabethes cyaneus*
SO: somata array of JO neurons
SSOs: self-sustained oscillations
TMDs: transmembrane domains
*To. yanbarensis*: *Topomyia yanbarensis*
*T. r. septentrionalis*: *Toxorhynchites rutilus septentrionalis*
*Ur. lowii*: *Uranotaenia lowii*
*W. smithii*: *Wyeomyia smithii*
WBF: Wing Beat Frequency
ZT: Zeitgeber.

## Acknowledgements

This study was financially supported by JST FOREST (No. JPMJFR2147 to AK), Tokai Pathways to Global Excellence (T-GEx), part of the MEXT Strategic Professional Development Program for Young Researchers (0121an0002 to MPS), MEXT KAKENHI Grant-in-Aid for Research Activity Start-up (No. JP22K15159 to MPS), JSPS Short-term fellowship (PE19013 to MPS), JSPS Invitational Fellowships for Research in Japan (Short-term) (S22091 to DFE), International Principal Investigator (PI) Invitation Program, Nagoya University, Japan (to DFE) and the Human Frontier Science Program Organization (No. RGP0033/2021 to AK).

## Author contributions

Conceptualization, Y.Y.J.X., Y.M.L., M.P.S., and A.K.; Methodology, Y.Y.J.X., Y.M.L., T.S.O., T.T.L., W.T.C., W.L., D.F.E. and M.P.S.; Investigation, Y.Y.J.X., Y.M.L., T.T.L., W.T.C., and M.P.S.; Resources, D.F.E., M.A., M.P.S. and A.K.; Writing, Y.Y.J.X., Y.M.L., D.F.E., M.A., M.P.S., and A.K.; Funding Acquisition, D.F.E., M.A., M.P.S. and A.K.; Supervision, M.A., M.P.S. and A.K.

## Declaration of interests

The authors declare no competing interests.

## STAR Methods

### Key resources table

**STAR Methods:** **Key resources table**
| REAGENT or RESOURCE | SOURCE | IDENTIFIER |
| --- | --- | --- |
| Antibodies |  |  |
| Mouse monoclonal anti-SYNORF1 3C11 | Developmental Studies Hybridoma Bank | AB_528479 |
| Rabbit Polyclonal anti-Octopamine | immusmol | IS1033 |
| STAINperfect™ immunostaining kit A | immusmol | SP-A-1000 |
| Monoclonal anti-cAMP (cyclic adenosine monophosphate) | Thermo Fisher | 500-9534 |
| DAPI 4',6-Diamidine-2'-phenylindole dihydrochloride | Roche Life Science Products | 28718-90-3 |
| Cy™3 AffiniPure™ Goat Anti-Horseradish Peroxidase | Jackson Immuno Research | AB_2338959 |
| Goat anti-mouse IgG Secondary Antibody, Alexa Fluor™ 488 | Thermo Fisher | #A-11029 |
| Experimental models: Organisms/strains |  |  |
| *Aedes aegypti* (Liverpool strain) |  | N/A |
| Software and algorithms |  |  |
| Audacity (version 3.0.3) | Audacity Team | https://www.audacityteam.org/ |
| Deeplabcut | A. and M.W. Mathis Labs<sup>51</sup> | [https://github.com/DeepLabCut](https://github.com/DeepLabCut/DeepLabCut) |
| OBS 29.1.2 | Open Broadcaster Software | https://obsproject.com/ |
| ImageJ (version 1.54g) | ImageJ<sup>52</sup> | https://imagej.net/ij/ |
| VibSoft (version 6.0) | Polytec | https://www.polytec.com |
| MEGA (version 12) | MEGA Team<sup>58</sup> | https://www.mega-software.net/ |
| R (version 4.3.1) | R Core Team<sup>53</sup> | https://www.R-project.org/ |
| Spike2 (version 10.08) | Cambridge Electronic Design | https://ced.co.uk/products/spkovicn |

### Resource availability

#### Lead contact

Further information on all experiments conducted as part of this report, in addition to requests for resources, can be requested from the lead contact, Matthew P. Su.

#### Materials availability

This study did not generate new unique reagents.

#### Data and code availability

- All original, unprocessed IHC data, as well as all processed RT-qPCR, sequencing and electrophysiology analysis data, used for this study has been deposited at Mendeley Data with accession doi: 10.17632/3b5z4tnrr5.1.

Raw electrophysiology and DLC data reported in this paper will be shared by the lead contact upon request.

- Additional information regarding analysis protocols/ data collection is available via contacting the Lead contact.

### Experimental model

#### Mosquito rearing

*Aedes aegypti* (Liverpool strain) mosquitoes were reared in a 12LJhr:12LJhr light–dark (LD) cycle at 26-28 °C and 60-70% relative humidity (RH). Adults were provided with a constant source of 10% glucose water. Horse blood feeding when necessary was conducted using an Orinno blood feeding system.

All mosquitoes used for experiments were aged between 4 and 9 days old. At least two days prior to all experiments, mosquitoes were transferred into light, temperature and humidity-controlled incubators set to 28°C and 60% RH. Light stimulus, provided by HIPARGEROLED (Hipargero) lights, followed a 12:12hr LD pattern.

### Method details

#### Immunohistochemistry

IHC followed previously published protocols^10,17^. Whole mosquito heads were removed on ice and fixed with 4% paraformaldehyde (PFA) in PBS (163-20145, Wako Pure Chemical Industries, Ltd) containing 0.25% Triton X-100 (PBT; #X100-500ML, Sigma-Aldrich) for 1 hr. Heads were then embedded in 6% Agarose (16520-050, invitogen) and the albuminAgarose blocks were solidified for 10 min at 4°C before fixing 4% PFA overnight at 4°C. Samples were washed in 100% methanol for 10 min, then washed in PBS (#T9181, Takara) for 30 min and sectioned (40 µm) using a vibratome (VT 1200S, Leica).

Sections were washed three times in 0.5% PBT and blocked in 10% normal goat serum (NGS; S-1000, funakoshi) / 0.5% PBT for 1 hr at room temperature. Primary antibodies in 10% NGS were then added and the samples stored overnight at 4°C.

After three washes with 0.5% PBT, samples were incubated with secondary antibodies in 10% NGS for 2 hr at room temperature. Following this, samples were washed three times with 0.5% PBT before a final PBS wash and mounting. Samples were imaged using a confocal microscope (FV3000, Olympus) equipped with a 60× Plan-Apochromat objective lens (UPlanSApo, NA = 1.3).

Octopamine IHC followed a protocol adapted from STAINperfect™ immunostaining kit A (SP-A-1000, immuno). Whole mosquito heads were removed on ice and fixed with 50% Fixation Reagent (SP-A-1005, immuno) in Fixation buffer (SP-A-1004, 1:25 in miliQ water, immuno) for 1hr. After removing Fixation Reagent, heads were washed 5 times by Wash Solution 1 (SP-A-1001, 1:25 in miliQ water, immuno) for 30 min at room temperature. Heads were then embedded in 6% Agarose and the Agarose blocks were solidified for 10 mins at 4°C before fixing 4% PFA overnight at 4°C. Samples were washed in 100% methanol for 10 mins, then washed in PBS for 30 mins and sectioned (40 µm) using a vibratome.

Sections were washed three times in Wash Solution 1 for 45 min and incubated in Permeabilization solution (SP-A-1006, 1:3 in Wash solution 1, immuno) for 20 min at room temperature. Samples were then washed three times in Wash Solution 1 for 45 mins and incubated in Stabilization reagent (SP-A-1008, 1 vial is reconstitute with 4 mL of Stabilization Buffer, immuno; SP-A-1008, 1:25 in miliQ water, immuno) for 1.5 hr at room temperature. After this, sections were washed three times in Wash Solution 1 for 45 mins and incubated in Saturation Solution (SP-A-1009, immuno) for 30 min at room temperature. Finally, sections were washed three times in Wash Solution 1 for 45 min and incubated in primary antibodies in Ab Diluent (SP-A-1010, immuno) overnight at 4°C.

After three washes with Wash Solution 1 for 45 mins at room temperature, samples were incubated with secondary antibodies in 10% NGS for 2 hr at 4°C overnight. Following this, samples were washed three times with Wash Solution 3 (SP-A-1003, 1:25 in miliQ water, immuno) before a final PBS wash and mounting. Samples were imaged using a confocal microscope (FV3000, Olympus) equipped with a 60× Plan-Apochromat objective lens (UPlanSApo, NA = 1.3).

Primary antibodies used included anti-octopamine (1:500, rabbit-Polyclonal, IS1033, immuno), cAMP (1:200, Thermo Fisher) and 3C11 (AB_528479, 1:30, Developmental Studies Hybridoma Bank (DSHB), University of Iowa). Secondary antibodies used included DAPI (1:5000, 28718-90-3, Sigma), Alexa Fluor Dyes (1:300, ThermoFisher) and Horseradish peroxidase (HRP; 1:100, Jackson Immuno Research).

#### RT-qPCR

Groups of virgin male or (non-blood fed) female mosquitoes were flash-frozen in liquid nitrogen and stored in a -80°C freezer. Mosquitoes were collected at Zeitgeber time [ZT] -12 of their entrainment regime. Tissues were then dissected on ice in RNAiso (#9109, Takara Bio Inc.), with pedicel (entire flagellum removed), flagellum, head (lacking both pedicels and mouthparts) and thorax (following leg and wing removal) tissue being collected.

Dissected samples were homogenized (Handheld Homogenizer, BT LabSystems) in RNAiso (#9109, Takara Bio Inc.). Additional RNAiso was added to the lysed samples to make up a final volume of 1 mL. The samples were then inverted several times before incubation at room temperature for 5 min. 200 µL chloroform (Kanto Chemical CO. INC.) was added to each sample and mixed well before incubating at room temperature for 15 min. Samples were then centrifuged at 12,000 x g for 15 min at 4°C before the addition of 0.5 mL 2-propanol (Sigma). Samples were stored in -20°C for 30 min followed by centrifugation at 12,000 x g for 10 min at 4°C, the supernatant was discarded and RNA pellet was washed with 1ml of freshly prepared 75% ethanol (Sigma) solution. Samples were centrifuged at 7,500 x g at 4°C for 5 min. The ethanol wash cycle was performed twice before the RNA pellet was dissolved in Nuclease Free Water (Invitrogen). RNA sample quality was confirmed using a Nanodrop and stored in -80°C until use.

RNA was reverse transcribed (ReverTra Ace^TM^qPCR RT Master Mix with gDNA Remover, FSQ-301, TOYOBO) prior to conducting qPCR (THUNDERBIRD^TM^SYBR^®^qPCR Mix, QPS-201, TOYOBO). For qPCR, a housekeeping gene *ribosomal protein S7* (*RPS7*) was used as the internal control. Octopamine receptor primers were designed using the NCBI primer design tool (Table S1). For each qPCR 96-well plate (Bio-Rad), three technical repeats for each sample and primer were tested, with the median of these technical values being used for analyses. In total, 8 repeats for male and 7 repeats for female were conducted for each tissue type.

#### Laser Doppler vibrometry & electrophysiology: Mosquito preparation

All recordings were conducted within the two hours prior to complete darkness (ZT11-13) at a temperature of 22±2°C.

Preparation of mosquitoes followed previously published protocols. Mosquitoes were sedated on ice, then glued to a small plastic rod. Glue (NORLAND PRODUCTS INC., 81) was minimally applied to the body to avoid hindering flagellar movement or obstructing major spiracles. After application of glue, only the right flagellum was free to move.

The rod was then held securely by a micromanipulator placed upon a vibration isolation table. Mosquitoes faced the laser directly and were positioned such that the flagellum was at a 90° angle to the vibrometer. For male mosquitoes, the laser focal point was chosen to be on the second flagellomere from the tip, whilst for females the third flagellomere from the tip was chosen.

#### Laser Doppler vibrometry & electrophysiology: Compound preparation

1 mM octopamine hydrochloride (OA; 770-05-8, Sigma), 1 mM clonidine (CD; 4205-91-8, Sigma), 1 mM phentolamine (PL; 65-28-1, Sigma),1 mM epinastine hydrochloride (EP; 108929-04-0, TCI) and 1 mM 8-Bromoadenosine 3′,5′-cyclic monophosphate sodium salt (8-Br-cAMP; 76939-46-3, Sigma) solutions were diluted from 10 mM stock solutions in Ringer solution, respectively. Mixed solution 1 mM clonidine plus BAPTA-AM (126150-97-8, TCI) in 1% DMSO in Ringer was made from 10 mM clonidine stock in Ringer and 1 M BAPTA-AM in 100% DMSO. Ringer solution alone was used for control injections^54,55^. Sharpened glass microcapillaries (G1; Narishige) were prepared using a puller (PC-10; Narishige), then filled with solution immediately prior to experiments.

#### Laser Doppler vibrometry & electrophysiology: recordings

Electrophysiology assays relied on the use of electrostatic actuation to provide sweep and step stimuli to mosquito flagellae. First, a reference electrode was inserted into the mosquito thorax to enable charging of the insect to -30V. A recording electrode was next inserted into the base of the mosquito JO. Copper actuators were then equipositioned either side of the flagellum to enable stimulus playback (with all stimuli generated using the Spike2 software), meanwhile, stimulated flagellum movement was measured by laser beam from laser Doppler vibrometry.

Looped sweep stimuli were applied to each individual mosquito. Each loop of sweep stimulation consists of 10 sets of sweeps, with each set comprising 4 different sweeps: forward phasic, forward anti-phasic, backward phasic and backward anti-phasic. Between phasic and anti-phasic sweeps of the same frequency orientation, a 0.1 s of silence was provided. A 0.4 s of silence was provided between sweep sets. A few step stimuli were conducted to each individual for nerve signal checking. Flagellum displacement and nerve recording data, as well as laser quality data, were recorded simultaneously using the Spike2 software 10.08 (Cambridge Electronic Design Limited).

After baseline recordings (pre-injection) of mechanical and electrical tuning, which were conducted for more than 5 min, a microcapillary for compound injection was inserted directly underneath the pedicel. Following injection, mechanical/electrophysiological recordings were then made for the next 15 min.

Sample sizes for each group:

Male Ringer injection = 14

Male 1 mM octopamine injection = 14

Male 1 mM epinastine injection = 14

Male 1 mM clonidine injection = 16

Male 1mM clonidine followed by 1mM epinastine injection = 4

Male 1 mM clonidine followed by 1mM phentolamine injection = 4

Male 1 mM clonidine plus BAPTA-AM injection = 4

Male 1 mM 8-Br-cAMP injection = 14 Female Ringer injection = 7

Female 1 mM octopamine injection = 7

#### Laser Doppler vibrometry & electrophysiology: analysis

For flagellar ear mechanical tuning analyses, a DC remove function with a time constant of 0.01 s was applied to the laser channel data, followed smoothing with a time constant of 0.0005 s to define the envelope of the response^12^. An Ultra-spline fit with span equals 0.1 in R was then applied, enabling identification of time point when the magnitude of the flagellar vibration reached its’ maximum (Fig S3). Calculation of the time at which has maximum flagellar vibration allowed for estimation of the stimulus frequency at that time, hereafter referred to as flagellar ear peak mechanical tuning frequency.

For electrical tuning analyses, sequential phasic and anti-phasic nerve responses were first averaged to cancel out artefacts recorded in the nerve channel due to electrostatic actuation (Fig S3). Firstly, a DC remove function with a time constant of 0.01s was applied to the nerve channel data. An Ultra-spline fit with span equals 0.5 for males and 0.8 for females, respectively, in R was then applied, enabling identification of time point when the magnitude of the action potential reached its’ maximum (Fig S3). Identification of the time at which has maximum magnitude of action potential facilitated calculation of the corresponding stimulus frequency, also referred to as ear peak electrical tuning frequency.

For the mechanical tuning frequency of an individual ear, the average peak frequency for each round of sweeps was taken from four sweeps (Forward phasic, forward anti-phasic, backward phasic and backward anti-phasic sweeps)^12^. A loess fit with span equals 0.1 was applied for data smoothing and the median value of average frequencies through first five minutes (pre-injection) was calculated as a baseline. This baseline was then subtracted from all mechanical tuning frequency estimates to calculate Δ Frequency. The maximum of Δ Frequency values after injection was taken as the max Δ Frequency per mosquito.

For electrical tuning frequency, the average between the forward and backward averaged sweeps were first computed. The same process as described above for the mechanical tuning frequency analysis was conducted to calculate electrical tuning frequency change Δ Frequency and max Δ Frequency per mosquito.

#### Fibrillae states: Recordings

Mosquitoes were prepared for fibrillae state recordings as described above for electrophysiology recordings. The position of fibrillae lining the mosquito’s flagellum was recorded using the camera built into the laser Doppler vibrometer equipped with a 10×Plan-Apochromat objective lens (M Plan Apo, NA = 0.28, Mitutoyo). Recordings lasted for 20 min (5 min before and 15 min after injection). Simultaneous measurement of unstimulated flagellar ear vibrations was conducted using Spike2.

Sample sizes for each group:

Male Ringer injection = 12

Male 1 mM octopamine injection = 14

Male 1 mM epinastine injection = 12

Male 1 mM clonidine injection = 12

Male 1 mM 8-Br-cAMP injection = 14

Female Ringer injection = 6

Female 1mM octopamine injection = 6

#### Fibrillae states: Analysis

All recorded videos were loaded into the Deeplabcut [DLC] learning system for frame extraction and labeling^51^. In total, the following three points of each flagellum were labeled as shown in Fig S5B.

Bottom (x,y) = Last segment on flagellum

BR (x,y) = Tip of hair locate on right last segment

BL (x,y) = Tip of hair locate on left last segment

A training dataset was then created including data from all recordings. Finally, the analyzed novel videos based on the evaluations enabled the creation of labeled videos via output filtered data.

The atan2 function in R software was used to calculate the angle based on the filtered data as follows:

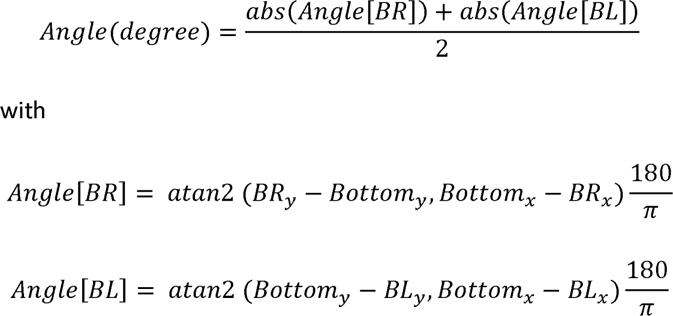

The loess fit in R then allowed for estimation of angle changes over time for each recording.

#### Unstimulated flagellum mechanical tuning: Analysis

The flagellum displacement recording in spike was output as TXT files and then loaded in R for further analysis. Recording data was split into multiple 1 s length segments and a Fast Fourier Transform (FFT) applied to each data segment. In a targeting frequency range (450 to 900 Hz for males and 150 to 400 Hz for females, respectively), the peak frequency of each segment was identified when displacement reached maximum. This frequency was taken as the peak mechanical tuning frequency at that timepoint.

#### Calculation of flagellar displacement change in ratio following injection

For each individual mosquito, the median peak displacement of pre-injection state was calculated and set as baseline (Disp0). Then peak displacement changes (Δ Disp) throughout 20 min recording was calculated by absolute displacement value minus baseline. Next, displacement changes were normalised by the baseline (Δ Disp/Disp0) and the ln of this ratio was then calculated [ln (Δ Disp/Disp0), shown as (ln Δ Disp ratio) in Fig 4D]. For frequency and displacement analysis of unstimulated fibrillae, values calculated during the injection time window were excluded because of off target effects induced because of the injection process.

#### RT-PCR

Groups of 5 mosquitoes were crushed in eppendorfs using pipette tips containing a buffer solution comprised of 10 mM Tris-Cl pH 8.2, 1 mM EDTA, 25 mM NaCl and 200 ug/ml Proteinase K. Samples were then incubated at room temperature for 30 min before being heated to 95°C for 2 minutes to inactivate Proteinase K.

RNA was reversed transcribed as described above for RT-qPCR experiments, then used to run PCR (Bio-Rad T100). PCR products were run on a 1% agarose gel using electrophoresis, including staining with DNA-staining fluorescent dye (WSE-7130 EzFluoroStain DNA, ATTO) and imaging using a Transilluminator (Daihan Scientific, WUV-M20). Bands were cloned out and submitted for sequencing using primers contained in Table S2 to cover different exons of both AAEL025638 and AAEL004245.

#### Analysis of Oct2αR receptors across species

Protein BLAST was used to identify the 5000 closest Dipteran orthologs of the amino acid sequence of *D. melanogaster* Octα2R (CG18208). Sequences for these orthologs were downloaded from NCBI and information associated with each gene were downloaded from NCBI or metazoa mart. MMseqs2 was used for protein sequence clustering, APcluster was used for visualization, while DEEPTMHMM and GPCRHMM were used for GPCR domain prediction^25–27,56^. TranslatorX was used to obtain the predicted proteins sequences from CDS.

To construct the phyogenetic tree, predicted amino acid sequences for the gene *catalase*^57^ for all species in the AGAP000606, AAEL025638 and AAEL004245 clusters were obtained from the National Centre for Biotechnology Information (Accession numbers included in Supplemental File 3). Using MEGA software ver 12^58^, These sequences were aligned using the CrustalW algorithm. The phylogenetic tree was then created using Maximum Likelihood analysis with 1,000-fold bootstrap replicates. Bootstrap values are displayed as numbers next to each branch node.

#### Identification of OAR positive cells from Ae. aegypti transcriptomic atlas

All maps included in Fig S9, S10 and S11 were generated from the online transcriptomic atlas database^45^.

To identify putative JO neurons, we filtered cells in the head (comprised of whole head tissue including antennae and mouthparts) by cell type (‘mechanosensory neurons’) and gene expression (‘*inactive* (AAEL020482)’). We then applied additional gene expression filters for *Oct*α*2R* and *Oct*β*2R* (AAEL025638*/* AAEL005945) to identify putative JO neurons also expressing specific octopamine receptors in both sexes.

To filter octopamine receptor positive cells in the flagellum (referred to as the ‘antennae’ in the online database), we filtered cells only by *Oct*α*2R* and *Oct*β*2R* (AAEL025638*/* AAEL005945) expression for both sexes.

The gene expression filtering is always greater than 1.

#### Statistical analysis

P < 0.05 (prior to correction) was set as the significance level for all statistical tests conducted. Shapiro-Wilk tests were used to test for normality.

For all RT-qPCR experiments, median RPS7 values were used to calculate DeltaCq (ΔCq) values for each primer. ANOVA tests were then used for each sex to test for significant differences in expression within each set of primers between different tissues.

To evaluate the significance of changes in flagellar tuning frequency after single injection, loess curves were fit to time series plots of extracted frequencies for individual mosquitoes. The maximum difference in flagellar frequency (Max Δ Frequency) was calculated by subtracting the median baseline flagellar frequency prior to injection from the maximum frequency after injection. Tukey’s tests and t tests were then run to check for significant differences between pre- and post-injection states for males and females, respectively.

To evaluate the significance of changes in electrical tuning frequency after single injection, loess curves were fit to time series plots of extracted frequencies for individual mosquitoes. The maximum difference in flagellar frequency (Max Δ Frequency) was calculated by subtracting the median baseline flagellar frequency prior to injection from the maximum frequency after injection. Tukey’s tests and t tests were then run to check for significant differences between pre- and post-injection states for males and females, respectively.

To evaluate the significance of angle changes in fibrillae states after single injection, loess curves were fit to time series plots of calculated angle for individual mosquitoes. The maximum difference in angle changes (Max Δ Angle) was calculated by subtracting the median baseline fibrillae angle prior to injection from the maximum fibrillae angle after injection. Pairwise wilcox tests and t tests were then run to check for significant differences between pre- and post-injection fibrillae states for males and females, respectively.

To evaluate the significance of changes in unstimulated flagellar tuning frequency after single injection, loess curves were fit to time series plots of extracted frequencies for individual mosquitoes. The maximum difference in unstimulated flagellar frequency (Max Δ Unsti Frequency) was calculated by subtracting the median baseline unstimulated flagellar frequency prior to injection from the maximum frequency after injection. Tukey’s test and t test were then run to check for significant differences between pre- and post-injection states for males and females, respectively.

To evaluate the significance of changes in correlation between unstimulated flagellar frequency change (Δ Unsti Frequency) and angle change (Δ Angle) after single injection the Pearson correlation coefficient was conducted to calculate the correlation of pre and post states. Wilcoxon Signed-Rank Tests were then run to check for significant differences between pre- and post-injection states within a sex.

To evaluate the significance of changes in correlation between log ratio displacement change (ln Δ Disp) and angle change (Δ Angle) after single injection the Pearson correlation coefficient was conducted to calculate the correlation of pre and post states. Paired t tests and Wilcoxon Signed-Rank Tests were then run to check for significant differences between pre- and post-injection states for males and females, respectively.

## Notes

### Competing Interest Statement

The authors have declared no competing interest.

