## Supplemental Text for "cAMP-related second messenger pathways modulate hearing function in *Aedes aegypti* mosquitoes"

**Supplementary Figure Legends:**

**Figure S1:** **Octopamine has some overlap with presynaptic sites in JO neuron somata in female *Ae. aegypti* pedicels.**

(A) Schematic diagrams of female ear: flagellum (top left), fibrillae (top right), pedicel (bottom, left) and JO (bottom right). Magenta and orange represent JO and flagellar neurons in the ear, respectively. Green and cyan dots represent octopamine and presynaptic sites, respectively. Ax, axons of JO neurons; BP, basal plate; C, cilia of JO neurons; P, prong; SO, somata array of JO neurons. A, anterior; M, medial.

(B) Location of presynaptic sites in female ear. Sections of female JO (right), JO close-up (middle) and flagellum (right) are shown. Anti-horseradish peroxidase (HRP; magenta) used as neuronal marker, anti-SYNORF1 (3C11; cyan) labels presynaptic sites. Scale bar = 20 μm.

(C) Location of octopamine in female ear. Sections of female JO (left), JO close-up (middle) and flagellum (right). DAPI (blue) labels nuclei, Anti-HRP (magenta) used as neuronal marker, anti-octopamine (green) shows octopamine localization. Scale bar = 20 μm.

(D) Individual channels of presynaptic sites staining in the ears of both sexes. From left to right: female anti-SYNORF1 (3C11; cyan) labels presynaptic sites, female anti-horseradish peroxidase (HRP; magenta) used as neuronal marker, male anti-SYNORF1 (3C11; cyan) labels presynaptic sites, male Anti-HRP (magenta) used as neuronal marker. Scale bar = 20 μm. Blue line selected region indicates somata of JO neurons, yellow line selected region indicates cilia of JO neurons.

(E) Individual channels of octopamine stanning in the ears of both sexes. From left to right: female anti-octopamine (green; octopamine localization), female Anti-HRP (magenta; neuronal marker), male anti-octopamine (green), male Anti-HRP (magenta). Scale bar = 20 μm. Region contained within blue lines indicates somata of JO neurons, region contained within yellow lines indicates cilia of JO neurons.

(F) Background signals resulting from secondary antibody staining. Sections of flagellum and JO of females (left) and males (right). Anti-HRP (magenta) was used as neuronal marker, anti-rabbit secondary antibody (green) shows background resulting from use of testing with secondary antibody alone. Scale bar = 20 μm.

**Figure S2: RT-qPCR based profiling of octopamine receptors across different tissues in male and female *Ae. aegypti***

(A) Delta Cq of each of the eight octopamine-family receptors in male flagellum (pink), head (blue), pedicel (purple) and thorax (red). Bar shows median of delta Cq values for all repeats. Individual points represent delta Cq of an individual repeat. *, p ≤ 0.05; ***, p ≤ 0.001; ANOVA. Eight repeats conducted for each tissue.

(B) Delta Cq of each of the eight octopamine-family receptors in female flagellum (pink), head (blue), pedicel (purple) and thorax (red). Bar shows median of delta Cq values for all repeats. Individual points represent delta Cq of an individual repeat. *, p ≤ 0.05; ANOVA. Seven repeats conducted for each tissue.

**Figure S3: Vibrometry/electrophysiology analysis paradigm and female vibrometry/electrophysiology data analysis**

(A) Diagram of sweep stimulus used for laser & electrophysiology recordings, as well as typical mechanical and electrical responses. Top row: phasic forward (1-1000Hz) sweep, anti-phasic forward sweep, phasic backward (1000-1Hz) sweep, anti-phasic backward sweep; middle row: laser channel recording; bottom row: nerve channel recording. Modified from Loh, Xu et al^2^.

(B) Laser data analysis. After a DC remove (time constant = 0.01) is applied to the raw data, it is rectified and smoothed (time constant = 0.0005). Modified from Loh, Xu et al^2^.

(C) Nerve data analysis. Data is duplicated, time shifted and then averaged to remove artefacts resulting from the use of electrostatic stimulation. Processed data is then analysed as for laser data. Modified from Loh, Xu et al^2^.

(D) Loess fits of mechanical (top) and electrical (bottom) tuning peak frequency changes from Ringer (control) and octopamine injection over 20 min in female *Ae. aegypti* mosquitoes. Ringer (green) and 1 mM octopamine (orange). Grey shaded region indicates before compound injection, colored regions after injection of compounds. Dark grey shaded area around loess fits indicates 95% confidence interval.

Sample sizes: Ringer injection = 7; octopamine injection = 7.

(E) Mechanical (top) and electrical (bottom) max Δ peak frequency change for Ringer (control) and octopamine injections. Middle line of boxplots represents median values of max mechanical and electrical peak frequency changes from compound injection. Each dot represents max peak frequency change of an individual mosquito. **, p ≤ 0.01; ***, p ≤ 0.001; t test.

**Figure S4: Representative images of changes in mechanical and electrical tuning following compound injection**

(A) Representative images of changes in mechanical tuning following compound injection for males (left four columns) and females (right column). Pre injection (black), Ringer (green), octopamine (orange), after octopamine effect is lost (light grey), clonidine (red), 8-Br-cAMP (yellow), epinastine (cyan), phentolamine (blue) and clonidine+BAPTA-AM (magenta).

(B) Representative images of changes in electrical tuning following compound injection for males (left four columns) and females (right column). Pre injection (black), Ringer (green), octopamine (orange), after octopamine effect is lost (light grey), clonidine (red), 8-Br-cAMP (yellow), epinastine (cyan), phentolamine (blue) and clonidine+BAPTA-AM (magenta).

**Figure S5: Workflow of Deeplabcut (DLC) data analysis and female fibrillae erection status/ unstimulated peak mechanical tuning analysis**

(A) Location of annulus in mosquito flagellae. Sections of male (left and middle) and female (right) flagellum. DAPI staining (blue) acts as nucleus marker. Male section with collapsed (left) and erect (middle) fibrillae show different conformational states of annulus (white circle), which appears absent in females (white circle). Scale bar = 5 μm.

(B) Workflow of Deeplabcut (DLC) for mosquito fibrillae movement quantification.

(C) Loess fits of female fibrillae angular movement following compound injection over 20 min. Ringer (green) and octopamine (orange). Grey shaded region indicates before compound injection, colored regions show after injection of compounds. Dark grey shaded area around loess fits indicates 95% confidence interval.

Sample sizes: Ringer injection = 6; OA injection = 6.

(D) Loess fits of female unstimulated mechanical tuning frequency change following compound injection over 20-min time period. Ringer (green), octopamine (orange). Grey shaded region indicates before compound injection, colored regions show after injection of compounds. Dark grey shaded area around loess fits indicates 95% confidence interval.

Sample sizes: Ringer injection = 6; OA injection = 6.

(E) Loess fits of female log ratio flagellar displacement change following compound injection over 20-min time period. Ringer (green), octopamine (orange). Grey shaded region indicates before compound injection, colored regions show after injection of compounds. Dark grey shaded area around loess fits indicates 95% confidence interval.

Sample sizes: Ringer injection = 6; OA injection = 6.

(F) Quantification of changes of maximum fibrillae angular movement (top) and maximum unstimulated mechanical tuning frequency change (bottom) following compound injection. Middle line of boxplots represents median value, whilst each dot represents data from an individual mosquito. ns, p > 0.05; **, p ≤ 0.01; t test.

**Figure S6: Correlation analysis of DLC and vibrometry data**

(A) Correlation between changes in peak delta unstimulated mechanical frequencies and fibrillae angles before and after compound injection for each injection type in males and females. Middle line of boxplots represents the median value, whilst each dot represents data from an individual mosquito. Dashed lines connect values for the same individual before and after injection. Ringer (green), octopamine (orange), clonidine (red), 8-Br-cAMP (yellow), epinastine (cyan). * p ≤ 0.05; Wilcoxon Signed-Rank Test.

(B) Correlation between log ratio change in unstimulated flagellar displacement and fibrillae angle changes before and after compound injection for each injection type in males and females. Middle line of boxplots represents median value, whilst each dot represents data from an individual mosquito. Dashed lines connect values for the same individual before and after injection. Ringer (green), octopamine (orange), clonidine (red), 8-Br-cAMP (yellow), epinastine (cyan). ns, p > 0.05; Pared t test for male, Wilcoxon Signed-Rank Test for female.

**Figure S7: Analysis of octα2R sequence data in *Aedes aegypti***

(A) Sequential locations of AAEL025638 and AAEL004245 on complementary strand of chromosome 1 of *Aedes aegypti* L5 genome.

(B) Outputs of DeepTMHMM analyses of AAEL025638 and AAEL004245. Four Transmembrane domains (TMDs) identified for AAEL025638 and three TMDs identified for AAEL004245.

(C) Amino acid sequence alignments for different Octα2Rs across species, including AAEL025638 and AAEL004245 from *Aedes aegypti*, AGAP000606 from *Anopheles gambiae* and FBGN0038653 from *Drosophila melanogaster*. Combined sequence of AAEL025638 and AAEL004245 also included.

(D) Sequencing using different primers (1L1 and C11) covering end of AAEL025638. Exon 3 of AAEL025638 not found in our sequencing data. CDS, Coding Sequence.

(E) Sequencing using different primers (1L1 and C11) covering start of AAEL004245. Exon 1 of AAEL004245 were not found in our sequencing data. CDS, Coding Sequence.

(F) Output of DeepTMHMM analysis of combined amino acid sequence of AAEL025638 and AAEL004245. Seven TMDs identified in total.

*A. aegypti*, *Aedes aegypti*; *An. gambiae*, *Anopheles gambiae*; *Drosophila melanogaster*, *D. melanogaster*.

**Figure S8: Analysis of octα2R sequence data across species**

(A) apcluster output showing clusters of different Octα2R genes across species. 4 clusters identified, centred on AAEL025638, AAEL004245, AGAP000606 and fbgn28653. Numbers indicate number of genes in each cluster. Representative *Ae. aegypti* Octα2R clusters centred on red (AAEL025638) and green (AAEL004245), representative *An. gambiae* Octα2R cluster centred on blue (AGAP000606), representative *D. melanogaster* Octα2R cluster centred on cyan (FBGN0038653).

(B) Amino acid sequence alignments for different Octα2Rs across all species in which two copies of Octα2R were identified.

(C) Phylogenetic tree (created based on the gene *catalase*) for all mosquito species identified in the AGAP000606, AAEL025638 and AAEL004245 clusters. Number next to each branch represents percentage of replicate trees where associated taxa clustered together. Number next to each species name represents the number of copies of Octα2R identified in that species. Branch length (scale bar: 0.05 units) represents number of amino acid substitutions for each sequence site. Magenta background covers species from tribe Aedini. Full list of accession numbers provided in Supplemental File 3.

(D) Amino acid sequence alignments for different Octα2Rs across all species in which two copies of Octα2R were identified showing end of AAEL025638 orthologs.

(E) Amino acid sequence alignments for different Octα2Rs across all species in which two copies of Octα2R were identified showing start of AAEL004245 orthologs.

(F) Output of DeepTMHMM analysis of combined amino acid sequence of AAEL025638 and AAEL004245 orthologs in *Aedes albopictus*. Seven Transmembrane domains (TMDs) identified but this combined sequence is still not classified as a GPCR by GPCRHMM.

*A. aegypti*, *Aedes aegypti*; *A. albopictus*, *Aedes albopictus*; *An. arabiensis, Anopheles arabiensis*; *An. coluzzii*, *Anopheles coluzzii*; *An. coustani*, *Anopheles coustani*; *An. darlingi*, *Anopheles darlingi*; *An. funestus*, *Anopheles funestus*; *An. gambiae*, *Anopheles gambiae*; *An. machlipalpis*, *Anopheles machlipalpis*; *An. marshallii*, *Anopheles marshallii*; *An. merus*, *Anopheles merus*; *An. moucheti*, *Anopheles moucheti*; *An. nili*, *Anopheles nili*; *An. stephensi*, *Anopheles stephensi*; *ASubRV.* *Armigeres subalbatus*; *Cx. p. pallens*, *Culex pipiens pallens*; *Cx. quinquefasciatus*, *Culex. quinquefasciatus*; *Drosophila melanogaster*, *D. melanogaster; Mal. genurostris*, *Malaya genurostris*; *Oc. camptorhynchus*, *Ochlerotatus camptorhynchus*; *Sa. cyaneus*, *Sabethes cyaneus*; *To. yanbarensis*, *Topomyia yanbarensis*; *T. r. septentrionalis*, *Toxorhynchites rutilus septentrionalis*; *Ur. lowii*, *Uranotaenia lowii; W. smithii*, *Wyeomyia smithii*.

**Figure S9: Analysis of single-nucleus RNA sequencing data of male *Aedes aegypti* head**

(A) Map of the cells annotated as male mechanosensory neurons (black) in the head.

(B) Map of male *iav* positive cells in cells annotated as mechanosensory neurons (purple) in the head.

(C) Map of male *iav* and octopamine alpha 2 receptor positive cells in cells annotated as mechanosensory neurons (cyan) in the head

(D) Map of male *iav* and octopamine beta 2 receptor positive cells in cells annotated as mechanosensory neurons (cyan) in the head.

*iav*, *inactive* (*AAEL020482*)

**Figure S10: Analysis of single-nucleus RNA sequencing data of female *Aedes aegypti* head**

(A) Map of the cells annotated as female mechanosensory neurons (black) in the head.

(B) Map of female *iav* positive cells in cells annotated as mechanosensory neurons (purple) in the head.

(C) Map of female *iav* and octopamine alpha 2 receptor positive cells in cells annotated as mechanosensory neurons (orange) in the head.

(D) Map of female *iav* and octopamine beta 2 receptor positive cells in cells annotated as mechanosensory neurons (orange) in the head.

*iav*, *inactive* (*AAEL020482*)

**Figure S11: Analysis of single-nucleus RNA sequencing data of male and female *Aedes aegypti* antennae/flagellum**

(A) Map of male octopamine alpha 2 receptor positive cells in cells annotated as mechanosensory neurons (left) and octopamine beta 2 positive cells in cells annotated as mechanosensory neurons (right) in the flagellum (registered as antennae). Circled region indicates mechanosensory neurons.

(B) Map of male *ppk317*_octopamine alpha 2 positive cells (left) and male *ppk317*_octopamine beta 2 positive cells (right) in the flagellum (registered as antennae). Circled region indicates *ppk317* positive cells. *ppk*, *pickpocket.*

(C) Map of female octopamine alpha 2 receptor positive cells in cells annotated as mechanosensory neurons (left) and octopamine beta 2 positive cells in cells annotated as mechanosensory neurons (right) in the flagellum (registered as antennae). Circled region indicates mechanosensory neurons.

**Supplemental Files**

**Supplemental File 1: Information on all identified Octα2R orthologs included in analyses shown in Figs S7 and S8**

**Supplemental File 2: GPCRHMM predictions for all identified Octα2R orthologs included in analyses shown in Figs S7 and S8**

**Supplemental File 3: List of accession numbers for the gene *catalase* for each species shown in phylogenetic tree in Fig S8C.**

**Supplementary Tables**

**Table S1: List of primers used for RT-qPCR**

| **Gene name** | **ensembl ID** | **Primer direction** | **Sequence** |
| --- | --- | --- | --- |
| Octα2R | AAEL004245 | Fwd | TAGGATCTACTACGCGGCGA |
|  |  | Bwd | CGCGTGAAACTTGTGTCCTG |
| Tyr1R | AAEL004396 | Fwd | AACCGCTCAACTACTCTCGC |
|  |  | Bwd | GTACCAGCCGAGAATGGGAG |
| Tyr2R | AAEL004398 | Fwd | GTCCGAAGCAGGGAACCTAC |
|  |  | Bwd | GAAGACCTCGGTGCTGTCAA |
| Octβ2R | AAEL005945 | Fwd | TCGTTCTTCATCCCCTGCAC |
|  |  | Bwd | CAACTGCTTCTCCTGTCGGT |
| Octβ1R | AAEL005952 | Fwd | GAGCCTAAAGCAGCACAACC |
|  |  | Bwd | GTCAAAGTCCGTGTCCGTGT |
| OctTyrR | AAEL006844 | Fwd | CATTGATTGCCGGTGTGTGG |
|  |  | Bwd | AATTGACACGGGAACTCGCT |
| OAMB | AAEL021100 | Fwd | TGGGGTCCTTCTACATCCCC |
|  |  | Bwd | GAAACGGGTGCCCATACCTT |
| Octα2R | AAEL025638 | Fwd | CTACCCGAGCGGATACACAC |
|  |  | Bwd | TGGATGTTCTTGAGCGCCTT |
| RPS7 | AAEL009496 | Fwd | TCAGTGTACAAGAAGCTGACCGGA |
|  |  | Bwd | TTCCGCGCGCGCTCACTTATTAGATT |

Fwd, forward; Bwd, backward

**Table S2: List of primers used for RT-PCR and sequencing**

| **Primer ID** | **Primer direction** | **Sequence** |
| --- | --- | --- |
| 25638_fwd | Fwd | GCTGGTGATCATAGCCATTG |
| 4245_bwd | Bwd | CATCACTTGAACAGGATCCGC |

**Table S3: List of mosquito species with one/two copies of octα2R**

| **Species name** | **Number of predicted Octα2R copies** |
| --- | --- |
| *Aedes aegypti* | Two |
| *Aedes albopictus* | Two |
| *Anopheles coustani* | Two |
| *Anopheles marshallii* | Two |
| *Armigeres subalbatus* | Two |
| *Culex pipiens pallens* | Two |
| *Ochlerotatus camptorhynchus* | Two |
| *Toxorhynchites rutilus septentrionalis* | Two |
| *Uranotaenia lowii* | Two |
| *Anopheles arabiensis* | One |
| *Anopheles coluzzii* | One |
| *Anopheles darlingi* | One |
| *Anopheles funestus* | One |
| *Anopheles gambiae* | One |
| *Anopheles maculipalpis* | One |
| *Anopheles merus* | One |
| *Anopheles moucheti* | One |
| *Anopheles nili* | One |
| *Anopheles stephensi* | One |
| *Malaya genurostris* | One |
| *Sabethes cyaneus* | One |
| *Topomyia yanbarensis* | One |
| *Wyeomyia smithii* | One |

**Table S4: Mosquito species whose fibrillae erection status has previously been described, in addition to associated genomic data on octα2R**

| **Species name** | **Fibrillae erection status** | **Paper Reference** | **octα2R copy number** |
| --- | --- | --- | --- |
| *Aedes aegypti* | Erect | Roth^S1^ | Two |
| *Aedes albopictus* | Erect | Loh, Xu et al^S2^ | Two |
| *Aedes caspius* | Collapsed/Erected | Nielsen and Nielsen^S3^ | No genomic information |
| *Aedes taeniorhynchus* | Collapsed/Erected | Clements^S4^ | No genomic information |
| *Aedes triseriatus* | Collapsed/Erected | Clements^S4^ | No genomic information |
| *Aedes vexans* | Collapsed/Erected | Nielsen^S5^ | No genomic information |
| *Anopheles balabacenis* | Collapsed/Erected | Nielsen^S5^ | No genomic information |
| *Anopheles freeborni* | Collapsed/Erected | Clements^S4^ | No genomic information |
| *Anopheles gambiae* | Collapsed/Erected | Georgiades et al^S6^ | One |
| *Anopheles maculatus* | Collapsed/Erected | Nielsen^S5^ | No genomic information |
| *Anopheles maculipennis* | Collapsed/Erected | Nutall and Shipley^S8^ | No genomic information |
| *Anopheles quadrimaculatus* | Collapsed/Erected | Roth^S1^ | No genomic information |
| *Anopheles stephensi* | Collapsed/Erected | Nielsen^S5^ | One |
| *Coquillettidia pertubans* | Collapsed/Erected | Clements^S4^ | No genomic information |
| *Culex pipiens* | Erect | Roth ^S1^ | Two |
| *Culex theileri* | Collapsed/Erected | Nielsen and Nielsen^S3^ | No genomic information |
| *Culiseta inornata* | Short hairs | Nielsen^S5^ | No genomic information |
| *Deinocerites cancer* | Short hairs | Nielsen^S5^ | No genomic information |
| *Mansonia perturbans* | Collapsed/Erected | Nielsen^S5^ | No genomic information |
| *Opifex fucus* | Short hairs | Nielsen^S5^ | No genomic information |
| *Phosphora confinnis* | Collapsed/Erected | Roth^S1^ | No genomic information |
| *Phosphora ferox* | Collapsed/Erected | [Nielsen](https://Nielsen12)^S5^ | No genomic information |
| *Toxorhynchites brevipalpis* | Erect | Nijhout^S8^ | No genomic information |
